# Megabarcoding reveals a tale of two very different dark taxa along the same elevational gradient

**DOI:** 10.1101/2024.04.29.591578

**Authors:** Wenya Pei, Vivian Feng, Aslak Kappel Hansen, Liping Yan, Amrita Srivathsan, Wentian Xu, Haoran Sun, Jun Yang, Xiaochen Zhang, Nan Yang, Qiuyue Yuan, Dong Zhang, Rudolf Meier

**Author notes:** **Correspondence** Dong Zhang, School of Ecology and Nature Conservation, Beijing Forestry University, Qinghua East Road 35, 10083 Beijing, China. Rudolf Meier, Center for Integrative Biodiversity Discovery, Leibniz Institute for Evolution and Biodiversity Science, Museum für Naturkunde, Invalidenstrasse 43, 10115 Berlin, Germany. These authors have contributed equally to this study (Wenya Pei, Vivian Feng).

## Abstract

Our planet is entering the sixth mass extinction without detailed spatio-temporal distribution and abundance information for most insect species. Particularly lacking is this information for dark taxa although they contain most animal species. This problem can be addressed with high-throughput single specimen barcoding (megabarcoding), but it remain unknown to what extent one dark taxon can be used as a proxy for another because of similar biodiversity patterns. This is here tested and we show that two Diptera taxa (Phoridae, Mycetophilidae) have very different spatio-temporal distributions across the same elevational gradient. We applied COI megabarcoding to samples collected every 14 days over three seasons at six elevations (800 to 1,800 m) in a temperate forest ecosystem in Northern China (Baihua Mountain Reserve). Elevation, precipitation, and temperature were used as environmental variables to understand the data obtained for 17,179 specimens representing 492 mOTUs of Phoridae and 2,281 specimens of 148 mOTUs of Mycetophilidae. Congruent is that the alpha diversity for both taxa increased in the summer and there is evidence for a mid-domain effect and phylogenetic clustering in high elevations. However, the extent and pattern of seasonal variation differed considerably. Phorid communities exhibited shifts between spring, summer and autumn seasons, but surprisingly little difference between elevations. In contrast, mycetophilid communities were structured along the elevational gradient and dependent on precipitation. Our study highlights the urgent need for obtaining detailed spatio-temporal information for the most diverse dark taxa, but also illustrates that these data can be readily obtained with megabarcoding.

## INTRODUCTION

“It was the best of times, it was the worst of times, it was the age of wisdom, it was the age of foolishness, it was the epoch of belief, it was the epoch of incredulity, it was the season of light, it was the season of darkness, it was the spring of hope, it was the winter of despair, we had everything before us, we had nothing before us, we were all going direct to Heaven, we were all going direct the other way…” Charles Dickens

A Tale of Two Cities was published over 150 years ago, but its opening paragraph is also an apt description of our current state of knowledge of insect biodiversity. We are living at the best of times because a large number of promising tools are becoming available, but we are also living at the worst of times because extinction is faster than our ability to gather enough information to understand and protect insect biodiversity. We are thus entering a season of darkness in that the sixth mass extinction is underway (Dirzo & Raven, 2003; Wake & Vredenburg, 2008; Pimm et al., 1995; Barnosky et al., 2011; Cardinale et al., 2012) with pollution, pesticide use, climate and land use change endangering the survival of many species (Crooks et al., 2017; Hautier et al., 2014; Chapin et al., 2000). Yet, despite the fact that insects are providing essential ecosystem services and contain more species and biomass than vertebrates, we are also still living in the age of foolishness in that comparatively little baseline data are available because too many decision makers still live in the epoch of belief that it is sufficient to study charismatic taxa (Titley, Snaddon, & Turner, 2017; Mammola et al., 2020). Indeed, insects remain neglected although the extinction risk for invertebrates can be higher than for vertebrate species (Hochkirch et al., 2023; Brondizio et al., 2019). The [only] spring of hope is that there is a growing awareness in the public consciousness and conservation organizations of the crucial roles of invertebrates in ecosystem functioning and services (Weisser & Siemann, 2008; McCary & Schmitz, 2021) so that there is hope that the epoch of incredulity will give way to a season of light. Particularly urgent is answering fundamental questions regarding regions/habitats with the highest species diversity, rate of species turnover, and areas with significant concentrations of small-ranged species (Wagner et al., 2021). In other words, we have to collect the kind of data for neglected insect taxa that have long been available for charismatic species and that have made them the target of most biodiversity studies.

Obtaining the relevant data is particularly important for so-called “dark taxa” that are hyper-abundant and diverse (Hartop et al., 2022) and can be defined as clades for which <10% of all species are described and the estimated diversity exceeds 1,000 species (Hartop et al., 2022). These taxa are so poorly known that even very basic diversity and distribution patterns are only revealed now. For example, Srivathsan et al. (2023) showed that surprisingly the same 20 families of insects dominate Malaise trap samples worldwide regardless of habitat and climate. Furthermore, most of these taxa face severe taxonomic neglect, with no indications of increased research activities in recent years (Srivathsan et al. 2023). This neglect is partially due to the fact that traditional species-level sorting with morphological tools for bulk samples is overwhelmed by the sheer quantity of specimens collected by standard traps. An alternative, metabarcoding, can be used to capture broad diversity patterns, but struggles with yielding abundance information and revealing the full diversity of dark taxa, because the specimens tend to be very small and are outcompeted in the genetic soup by larger common species (see data in Iwaszkiewicz-Eggebrecht et al., 2023) although size sorting of bulk samples can remedy these problems for some species (Elbrecht et al., 2017). It remains, however, unknown whether the numerous rare taxa can be detected given that dark taxa follow the typical species abundance distributions (Callaghan et al., 2023) in that only few species are common, and most rare (Srivathsan et al., 2019; Hartop et al., 2022).

Fortunately, we are living in an age of rapid technological change, so that techniques are becoming available that allow for carrying out detailed specimen-based studies of insect communities at scales previously unheard of (D’Souza & Hebert, 2018; Braukmann et al., 2019; Yeo et al., 2021). In particular, “megabarcoding” (Chua et al., 2023), i.e., efficient DNA barcoding workflows applied to tens of thousands individual specimens, has made studying hyperdiverse taxa at species-level, with abundance data, manageable (Meier et al., 2016; Srivathsan et al., 2021; Yeo et al., 2020). The application of megabarcoding to all specimens in a sample can yield diversity and abundance information at molecular operational taxonomic unit (mOTU) level even for those mOTUs that consist of small, rare specimens. Given that the congruence between mOTUs and morphospecies boundaries tend to be high, megabarcoding data can be used as if mOTUs were species-equivalents (Wang et al., 2020). This allows for the analysis of community structure in so much detail that it occasionally more than rivals the data available for vertebrate clades because specimens and species are collected in far greater quantities.

We here study phorid flies (Diptera: Phoridae) and fungus gnats (Diptera: Mycetophilidae) that are classic examples of hyperdiverse and understudied dark taxa (Disney, 1994; Chimeno et al., 2022, 2023; Kurina & Grootaert, 2016). Srivathsan et al. (2023) found that both families ranked among the top 20 most species-rich insect families captured in Malaise traps globally. Phoridae are globally distributed encompassing only approximately 4,000 described species (Disney, 1994; Srivathsan et al., 2019) although the actual species diversity is potentially two orders of magnitude greater (Srivathsan et al., 2019). The larvae of this family demonstrate a broader range of habits than any other family of animals on Earth and develop and occupy a tremendous variety of microhabitats (Disney, 1990, 1994). Given their ubiquity, abundance, and diversity, phorids hold significant potential as a taxonomic group for environmental assessments (Disney & Durska, 2008). Despite this potential, previous ecological studies utilizing phorids for environmental assessments (Durska, 2013, 2020) have been limited in scope due to the challenge of processing a large number of specimens and the difficulty in their morphological identification. The second target clade is Mycetophilidae which is the most species-rich group of Diptera associated with fungi (Jakovlev, 2012). At the same time, mycetophilids are among the most abundant and diverse insect groups found in temperate and tropical forests (Ševčík, Hippa, & Burdíková, 2022). The family is the second most species-rich family of the suborder Bibionomorpha, with 233 genera and about 4,500 species, described from all biogeographic regions (Pape et al., 2011; Ševčík, Hippa, & Wahab, 2014). Mycetophilid adults are thought to be sensitive to drought and mainly inhabit humid and shaded forests, with preference for stable old-growth stands (Økland, 1994), thus making them excellent indicators in the assessment of forest habitats for nature conservation (Kallweit, 2013). They are believed to have low dispersal capabilities and are therefore likely vulnerable to environmental change (Jakovlev, 2012).

We here study the spatio-temporal diversity of these two dark taxa along an elevational gradient in the Baihua Mountain Reserve, a temperate forest ecosystem in China renowned for its high biodiversity, environmental heterogeneity, and diverse climate (Zhang et al., 2013; Pei et al., 2021). China is not only a megadiverse country, but also one of the countries with the most threatened biodiversity (Wang et al., 2020). Unfortunately, there are comparatively few quantitative studies investigating the biodiversity of non-charismatic insects and their distribution patterns in China, especially in temperate montane forest ecosystems (Zou et al., 2014; Jiang, 2006; Axmacher et al., 2011). Especially, the dark taxa have been neglected because morphological techniques are of limited use for processing very large numbers of specimens. For example, currently only 226 species of Phoridae and 260 species of Mycetophilidae have been documented in all of China (Liu et al., 2023; Ye et al., 2023) and we will here show that the number of phorid species in the Baihua Mountain Reserve greatly exceed that. Not surprisingly, spatio-temporal distribution data for these taxa are essentially non-existent.

We here use megabarcoding to test whether the spatio-temporal distribution patterns of two dark taxa resemble each other when studied from multiple perspectives. In particular, we use environmental data (precipitation, temperature) to understand how they affect spatio-temporal species diversity and turnover given that precipitation and temperature regimes are likely affected by global changes in climate. To test whether the results are robust to changes in temporal resolution and species abundance distributions, we analyze 2 and 4-week sampling intervals and vary the degree to which rare species were removed. This leads to recommendations which species are potential conservation targets and could be good indicators for environmental change. Our study thus not only generated barcode data for the Chinese fauna of two dark taxa, but also individual voucher specimens for further taxonomic work. Together, the barcodes and vouchers can serve as baseline data for future monitoring of phorid and mycetophilid flies by metabarcoding bulk insect samples or sequencing environmental DNA (eDNA) from ethanol.

## MATERIALS AND METHODS

### 2.1 Study site

The Baihua Mountain Reserve, situated at coordinates E115° 25′-115°42, N39° 48′-40 ° 05 ′, is located at the northern end of the Taihang mountain range in Beijing municipality, China, roughly 120 km west from the city center (Figure 1). It is one of the two national nature reserves close to Beijing’s city center (Zhang et al., 2013). Serving as a crucial ecological buffer for the city, the reserve lies in a temperate, semi-humid climate with continental characteristics, primarily influenced by seasonal wind patterns. Spring (March-May) is marked by notable drought conditions. Summer (June-August) experiences high temperatures and abundant precipitation. Autumn (September-November) offers a moderate transition from warm to cold, with reduced precipitation.

**FIGURE 1.**
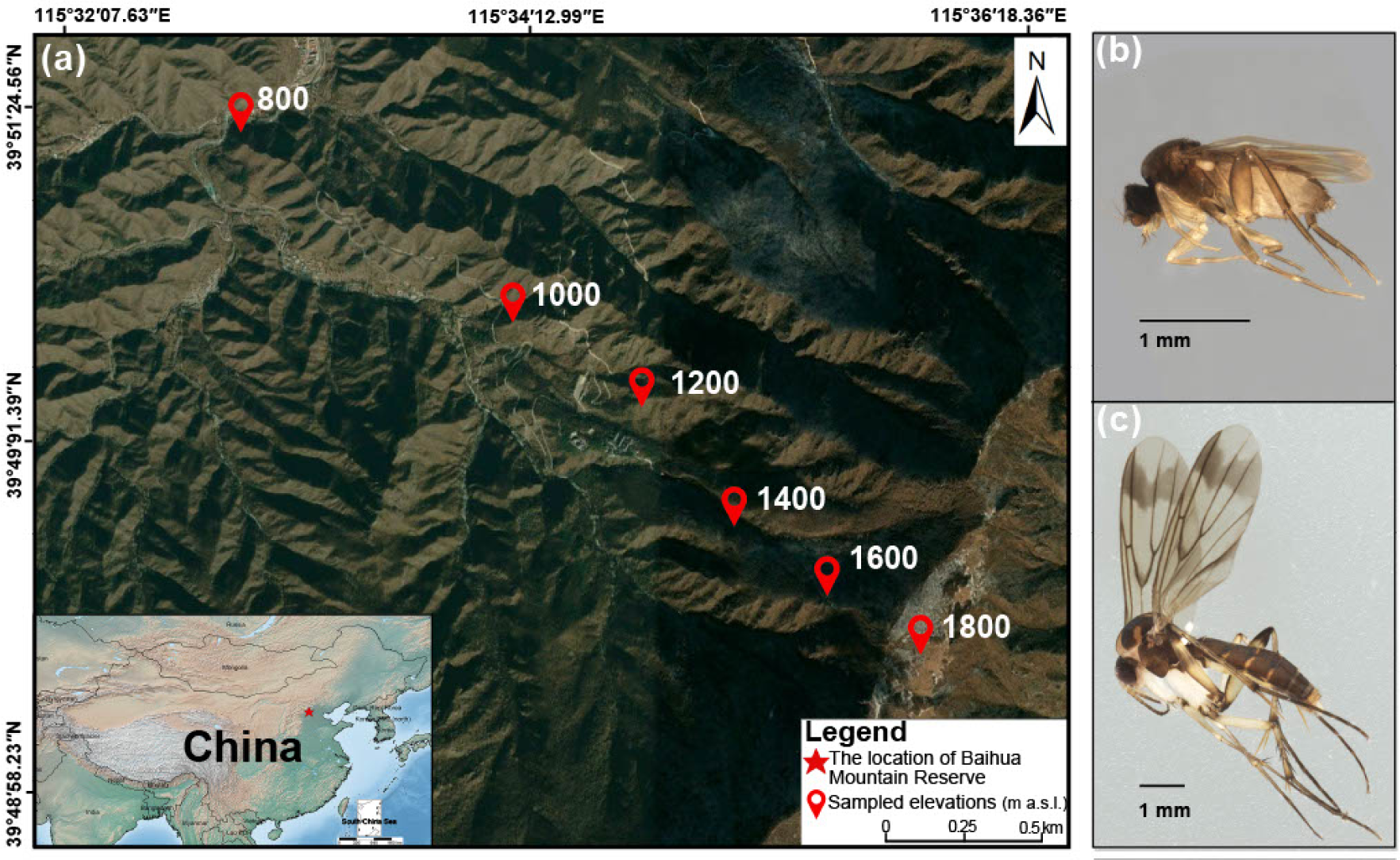
(a) Sampling localities in the Baihua Mountain Reserve, Beijing, China. (b) & (c) show the lateral habitus of the target two taxa, Phoridae and Mycetophilidae, respectively. The two specimens photographed were from the most abundant mOTUs of Phoridae and Mycetophilidae captured in this project.

### 2.2 Sampling along an elevational gradient

Samples were collected weekly using Townes-type Malaise traps (Townes, 1972) deployed at six sampling elevations (800, 1,000, 1,200, 1,400, 1,600 and 1,800 m a.s.l., Figure 1, Table S1) along a linear elevational transect ranging from the entrance of Baihua Mountain Reserve up to the mountain meadow. Sampling covered 37 active weeks (March-November 2019) yielding a total of 222 samples that were preserved in 95% ethanol. Samples were taken to the Museum of Beijing Forestry University and stored at -20 °C until DNA extraction.

### 2.3 Climate data

High spatial resolution (0.5’, ∼1 km) datasets for monthly mean temperature and monthly precipitation totals for all elevations in the Baihua Mountain Reserve were extracted from WorldClim 2.1 (Fick & Hijmans, 2017) based on latitudinal and longitudinal information using the ‘geodata’ (Hijmans et al. 2023) R (R Core Team, 2022) software package and are the average for the years 1970-2000.

### 2.4 Pre-sorting of specimens to putative species (mOTUs) with NGS barcoding

After initial sorting of individuals to the Diptera families Phoridae and Mycetophilidae, all specimens were processed following a similar approach to Srivathsan et al. (2021). DNA was extracted using 10 to 20 μl of HotSHOT lysis buffer (Truett et al., 2000) per specimen depending on size and heated to 65°C for 18 min, followed by 98°C for 2 min, followed by addition of an equal volume of neutralization buffer. A 658 bp fragment of cytochrome c oxidase subunit I (COI) was amplified using primers LCO1490: 5 ′ -GGTCAACAAATCATAAAGATATTGG-3 ′ (Folmer et al. 1994) and jgHCO2198: 5 ′ -TANACYTCNGGRTGNCCRAARAAYCA-3′ (Geller et al 2013). The primers used were labelled with 13 bp tags at the 5′ end designed for MinION-based barcoding (Srivathsan et al., 2021). Each PCR reaction contained 7 μl Mastermix (CWBIO 2xTaq MasterMix, CW0682), 1 μl bovine serum albumin (1 mg/ml), 1 μl of each primer (10 μM) and 4 μl of template DNA, for a total of 14 μl. The cycling conditions were as follows: initial denaturation at 95 °C (5 min), followed by 35 cycles of denaturation at 94 °C (1 min), annealing at 45 °C (2 min) and extension at 72 °C (1 min), succeeded by final extension of 72 °C (5 min). The PCR products were pooled in equal volumes and cleaned using 0.6X AMPure XP magnetic beads (Beckman Coulter). The DNA library was prepared with the Nanopore ligation kit (SQK-LSK114) and the NEBNext MinION Companion Module (E7180) and the products were sequenced on FLO-MIN114 R10.4.1 flowcells using the Oxford Nanopore Technologies Mk1B. Basecalling was conducted locally using Guppy (version 6.4.2) under the super accuracy model. Demultiplexing and barcode calling was done using ONTbarcoder (version 0.1.9) (Srivathsan et al., 2021) under default settings.

The barcodes were then matched to the NCBI nt-database using BLASTN in the BLAST+ suite (Camacho et al., 2009) with minimum e-value of 1e-5 and percentage identity match of 80%. The low threshold was chosen in consideration of the relatively unknown taxa of interest, which were sampled for the first time from this region. BLASTN matches were parsed using readsidentifier (Srivathsan et al., 2015) and barcodes which matched non-target taxa with a >90% similarity score were filtered out. Barcodes were aligned using MAFFT v7 (Katoh & Standley, 2013) with default parameters. To delineate putative mOTUs, we employed objective clustering through a Python script (https://github.com/asrivathsan/obj_cluster) that implements the objective clustering algorithm described by Meier et al. (2006). We obtained results for three p-distance thresholds (2, 3 and 4%), which encompass thresholds commonly employed in the literature for species delimitation (Ratnasingham & Hebert, 2013). Note that other species delimitation algorithms could have been used, but they generally yield very similar results (Yeo et al., 2021; Meier et al., 2022). Specimens yielding barcodes with low percentage identity match (<97%) to the NCBI database were verified to family by taxonomic experts (for Mycetophilidae) and the authors (for Phoridae).

### 2.5 Diversity analyses

Samples were grouped by elevation, at both 2 and 4-week intervals, giving two different datasets for each family (Tables S2-S5). To test the robustness of the results, some analyses were repeated with trimmed datasets from which varying degrees of rare species were removed (1: singletons removed, 2: single and doubletons removed, 3: mOTUs with less than 5 specimens removed, 4: mOTUs with less than 10 specimens removed). Community matrices were generated with a Python script (clusterlist_to_commatrix.py: see (Yeo et al., 2021)) which uses the objective clustering outputs from above. All statistical analyses were performed in R v. 4.2.0 (R Core Team, 2022) and all figures were produced with the ggplot2 package v. 3.4.2 (Wickham, 2016).

To assess the species richness at each elevation and month, samples were rarefied with iNEXT (Hsieh, Ma, & Chao, 2016; Chao et al., 2014) in R using 1,000 bootstrap replicates to account for unequal sampling completeness. We utilized sampling coverage as a proxy for gauging completeness. Subsequently, we tested for correlation between richness and abundance using the Pearson correlation coefficient function “cor()” in R for both datasets. This was also done for monthly total precipitation and monthly mean temperature. To compare patterns of richness with climate data, we used the ggpmisc package (Aphalo, 2023) to show equation of lm model and poly model, and model performances were evaluated by R-square values and p-value.

To study changes in richness along the elevation gradient and over time, we determined richness over two weeks for each elevation gradient, and then calculated the mean elevation per taxa at 2-week intervals (e.g. (Elevation A x number of mOTUs at elevation A for 2 weeks + Elevation B x number of mOTUs at elevation B for 2 weeks) / Total number of mOTUs at elevations A&B for 2 weeks) (Thorne et al., 2022).

To track shifts in elevation ranges for species between months, we compared elevational ranges month-by-month for each mOTU. For this purpose, we used three measures for each mOTU in each month: (a) highest elevation, (b) lowest elevation and (c) weighted mean elevation (average of elevations for all individuals of a given mOTU) (Menéndez et al., 2014). However, month-by-month elevational change can only be studied for mOTUs that appear in consecutive months (singletons and mOTUs appearing only in a single month are naturally excluded). We first identified the Phoridae and Mycetophilidae mOTUs which fulfilled the criteria of having at least one specimen appearing in two consecutive months. We then measured changes in elevation between sets of two consecutive months and tested for significance using nonparametric Wilcoxon signed - rank tests. Next, to ensure that observations for Phoridae held true even if scaled to the same richness and abundance as Mycetophilidae, we subsampled the month with the largest number of Phoridae mOTUs (August) in two different ways in order to match the number of mOTUs and the number of specimens of Mycetophilidae in August using the sample() function in R. Lastly, the analysis was repeated on the trimmed datasets.

For the analysis of community composition, we first excluded any samples (grouped by month and elevation) with <10 specimens. We then used Non-metric Multi-dimensional Scaling (NMDS) based on Bray-Curtis dissimilarity matrices to visualize the differences in community composition. The NMDS1 and NMDS2 coordinates were derived with the metaMDS function from the vegan package (Oksanen et al., 2022). This was repeated with each of the four trimmed datasets. Due to the much larger size of the Phoridae dataset, we repeated the analysis on subsamples matching the scale of the Mycetophilidae dataset to check if observed patterns still hold true. Subsampling was conducted in the same manner as above.

We tested for significant differences in community composition between elevations and between months separately with sequential permutational analyses of variance (PERMANOVA, 9,999 permutations) tests with the adonis2 function (Zhu et al., 2021).

We used the UpSetR package (Conway et al., 2017) based on the ‘UpSet’ technique (Lex et al., 2014; Lex & Gehlenborg, 2014) to visualize unique and shared mOTUs between different months and elevations for Phoridae and Mycetophilidae.

We used the beta.multi.abund and beta.pair.abund functions from the betapart package (Baselga & Orme, 2012) to obtain both multiple-site and pairwise dissimilarity values. We used β_BC.BAL_ and β_BC.GRA_ (Baselga, 2017) values to examine the proportion of overall dissimilarity explained by spatial turnover and nestedness respectively. This was repeated with the four trimmed datasets.

Our observed richness across elevations was compared against a predicted pattern following the Mid-Domain Effect (MDE) model given by rangemodelR package (Marathe, 2019), using the following parameters: 200 m for the elevation band interval, ‘sh’ for nature of the boundaries being soft at our lowest elevation and hard at our highest elevation (as the mountain range is not taller than 1,900 m a.s.l. in the study site), and 5,000 repetitions.

For both Phoridae and Mycetophilidae a separate MAFFT alignment (Katoh & Standley, 2013) containing a single center barcode for each mOTU was generated. For these, maximum likelihood trees were generated using IQtree (Minh et al., 2020) with partitioning by codon position and model search using ModelFinder (Kalyaanamoorthy et al., 2017). We quantified the mean-nearest-taxon-distance (MNTD) and the nearest-taxon-index (NTI) using the picante package (Kembel et al., 2010) to characterize phylogenetic community composition of the two taxa both along elevations and across months.

## RESULTS

### 3.1 Species delimitation based on NGS barcodes

We obtained 19,460 barcodes in total, 17,179 for Phoridae and 2,281 for Mycetophilidae (Table S6). For Phoridae, 2% p-distance gave 537 mOTUs, 3% p-distance gave 492 mOTUs and 4% p-distance gave 451 mOTUs. For Mycetophilidae, 2% p-distance gave 151 mOTUs, 3% p-distance gave 148 mOTUs and 4% p-distance gave 144 mOTUs. Most mOTU boundaries remained consistent, with variations of less than 9% across species delimitation parameters. For subsequent analyses, we used the mOTUs obtained at 3% p-distance.

### 3.2 Alpha-diversity

We found high sampling coverage for both taxa at most elevations and months thus justifying the use of richness in subsequent analyses (Tables S7 and S8). Note that high sampling coverage was obtained despite a positive correlation between richness and abundance for both groups (Phoridae: r = 0.7, n = 50, p < 0.001; Mycetophilidae: r = 0.92, n = 31, p < 0.001).

### 3.3 Changes in richness over time and space

A unimodal peak of alpha-diversity in August is observed for both taxa (Figure 2). Similarly, the months with highest abundance are driven by a few very abundant mOTUs (Tables S2 and S3). For Phoridae, eight very abundant mOTUs contribute 1,818 specimens resulting in the month of June having the highest overall abundance (4,024 specimens) (Table S2). Similarly, four Mycetophilidae mOTUs occurred in high numbers (558 specimens in total) in August, resulting in a clear abundance peak for this month (1,277 specimens) (Table S3). But there are also major differences between the taxa. Phoridae appeared earlier and disappeared later in the season than Mycetophilidae. Phoridae emerged in large numbers in early spring (March), especially at lower elevations where richness was highest. 40 mOTUs were still captured in November, and 14 mOTUs were captured at four elevations in late November (Table S4). In contrast, Mycetophilidae were not observed until the end of May, with only two singleton mOTUs recorded at lower elevations (800 m and 1,200 m a.s.l.), while the last mOTU was collected in November at the lowest elevation (800 m a.s.l.) (Table S5). The abundance was highest for Phoridae in June, but August for Mycetophilidae.

**FIGURE 2.**
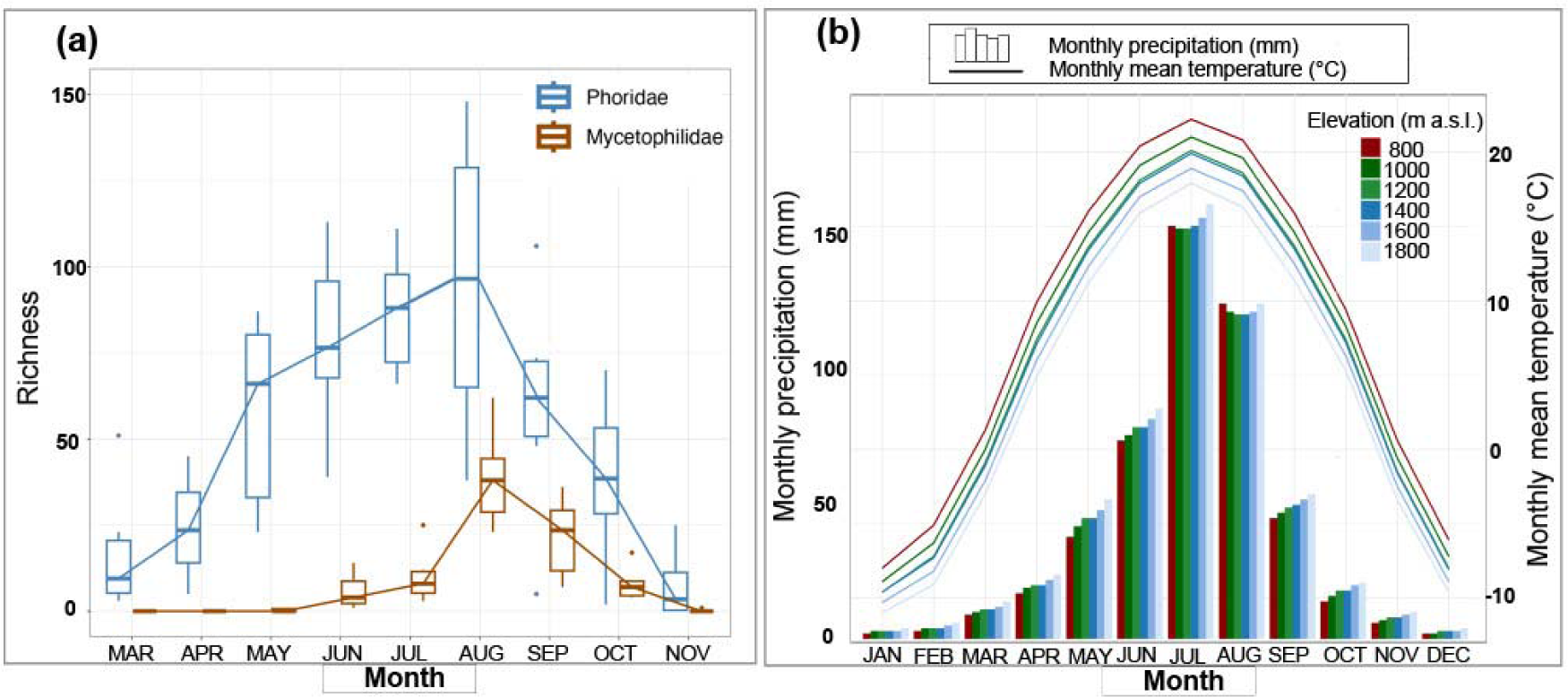
Variations in richness and climatic variables across months. (a) Richness and (b) total precipitation and mean temperature at six elevations in Baihua Mountain Reserve are strongly correlated (r = 0.83, n = 35, p < 0.001).

In spring, the richness of Phoridae along the elevation gradient showed a shift from low elevation to high elevation (800 m to 1,200 m a.s.l.) over time (Figure 3a). From spring to summer, the mean elevations of Phoridae show a noticeable upward trend along the elevation gradient. In summer, Phoridae communities showed a stable distribution pattern along the elevation gradient over time, showing a bimodal pattern with richness peaks at 1,200 m and 1,600 m a.s.l. Conversely, from summer to autumn, there is a shift observed in Phoridae distribution, moving from high elevations to lower elevations (Figure 3a). For Mycetophilidae, only two singleton mOTUs appeared in early spring, so that there are no discernible changes in elevation between spring and summer, but a shift downward can be seen from summer to autumn (Figure 3b). Note that Mycetophilidae exhibit limited activity until late July/early August. During this period, their activity is predominantly concentrated at lower elevations, gradually shifting to higher elevations throughout August (Figure 3b). In addition, with the arrival of the dry autumn, the number of Mycetophilidae mOTUs suddenly decreases which coincides with a shift to lower elevations (Figure 3b). These results are stable even when the trimmed datasets are used (Figure S1).

**FIGURE 3.**
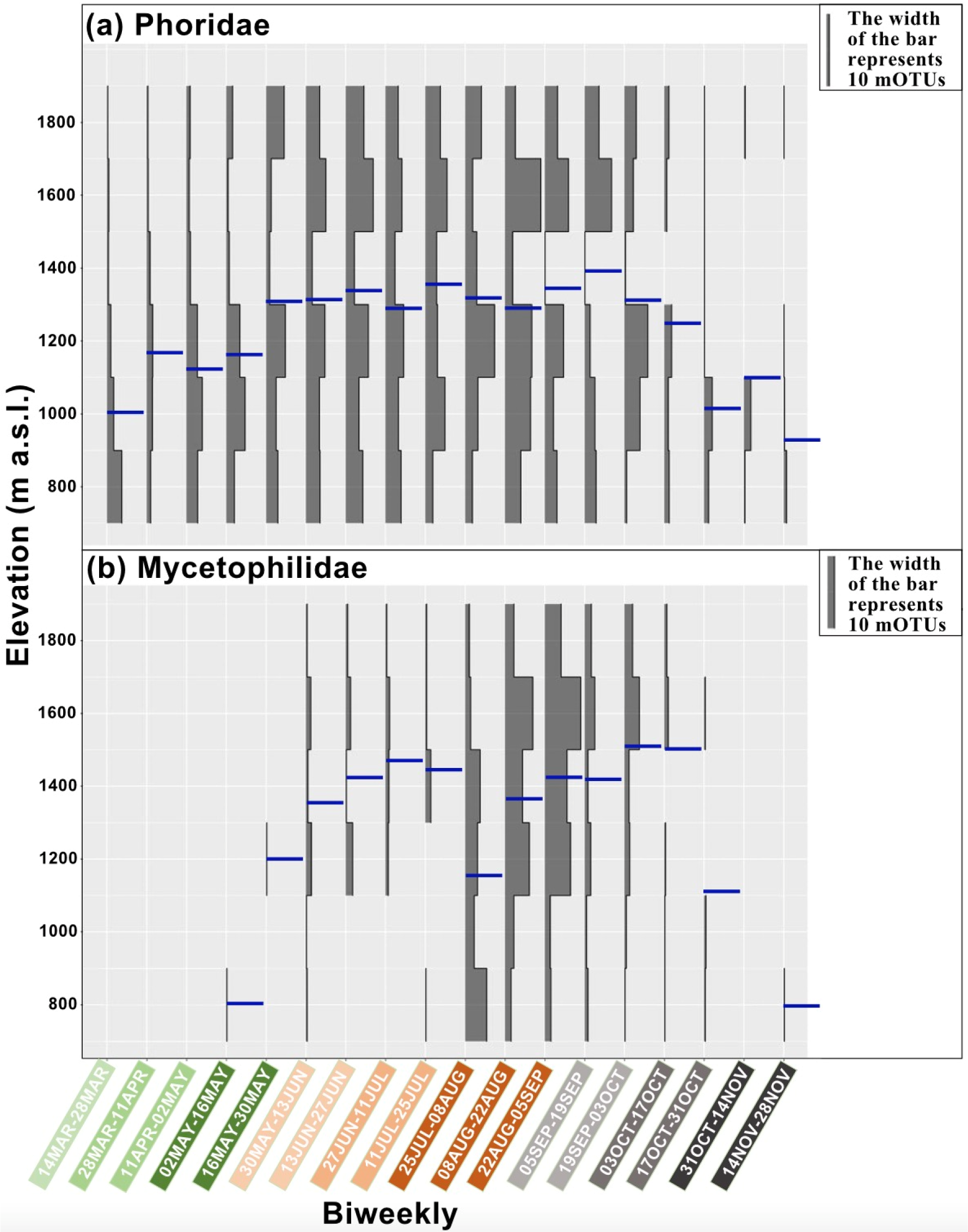
Patterns of richness along the elevation gradient, shown in 2-week intervals. Blue bar indicates mean elevation per taxa at 2-week intervals (see Methods for details).

### 3.4 Relationship between richness and environmental factors

Monthly mean temperature (r = 0.73, n = 50, p < 0.001) and monthly total precipitation (r = 0.69, n = 50, p < 0.001) have a significant relationship with richness of Phoridae and the fitted positive linear regressions (3.63 and 0.516 respectively) explain 54% and 47% of the observed variability, respectively (Figure 4a). For Mycetophilidae we instead found that the monthly total precipitation of the previous month was a better predictor of their richness (r = 0.67, n = 52, p < 0.001) than that the monthly total precipitation for the month (r = 0.28, n = 64, p = 0.02) and the fitted positive linear regressions (0.196) explain 46% of the observed variability, indicating a possible lag effect for richness following sometime after precipitation (Figure 4b).

**FIGURE 4.**
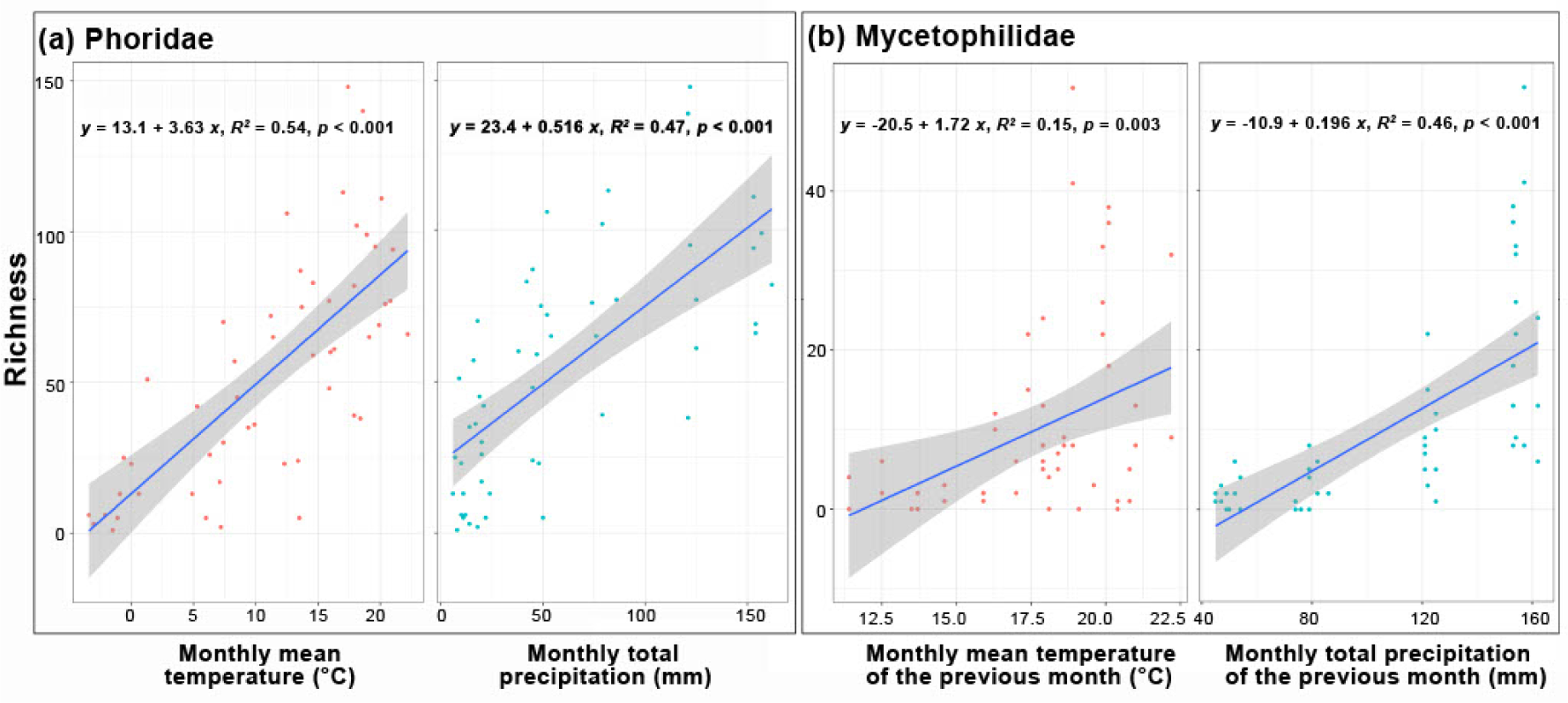
Linear correlation between richness of (a) Phoridae with monthly mean temperature and monthly total precipitation and (b) Mycetophilidae with the monthly mean temperature of the previous month and monthly total precipitation of the previous month.

### 3.5 Seasonal changes of elevation ranges

Significant seasonal elevation shifts were only detected for Phoridae mOTUs (see Figure 5c; Table S9). Phoridae exhibited a significant increase in elevation throughout spring and early summer, with increasing weighted mean elevation (March to April: V = 335.5, mean with SD = 116.39 ± 195.45, p = 0.003, April to May: V = 366, mean with SD = 125.57 ± 218.24, p < 0.001, May to June: V = 3677.5, mean with SD = 118.51 ± 277.48, p < 0.001) (Figure 5c, Table S9). No significant changes in elevation were observed in summer (June to August) or in the transition from summer to autumn (August to September). In autumn, a significant decrease in the weighted mean elevation was observed (September to October: V = 917, mean with SD = -135.96 ± 281.09, p < 0.001). The significance remained unchanged even for the trimmed datasets (Figures S2-S4, Table S9). In comparison, Mycetophilidae did not show significant changes in the weighted mean elevation during summer and autumn (June to October) (Figure 5c, Table S9). However, there was a significant increase in the weighted mean elevation from June to July for each of the trimmed datasets (no doubletons: V = 51, mean with SD = 134.08 ± 184.93, p = 0.019, no mOTUs <5 specimens: V = 51, mean with SD = 134.08 ± 184.93, p = 0.019, no mOTUs <10 specimens: V = 51, mean with SD = 146.27 ± 188.84, p = 0.019).

**FIGURE 5.**
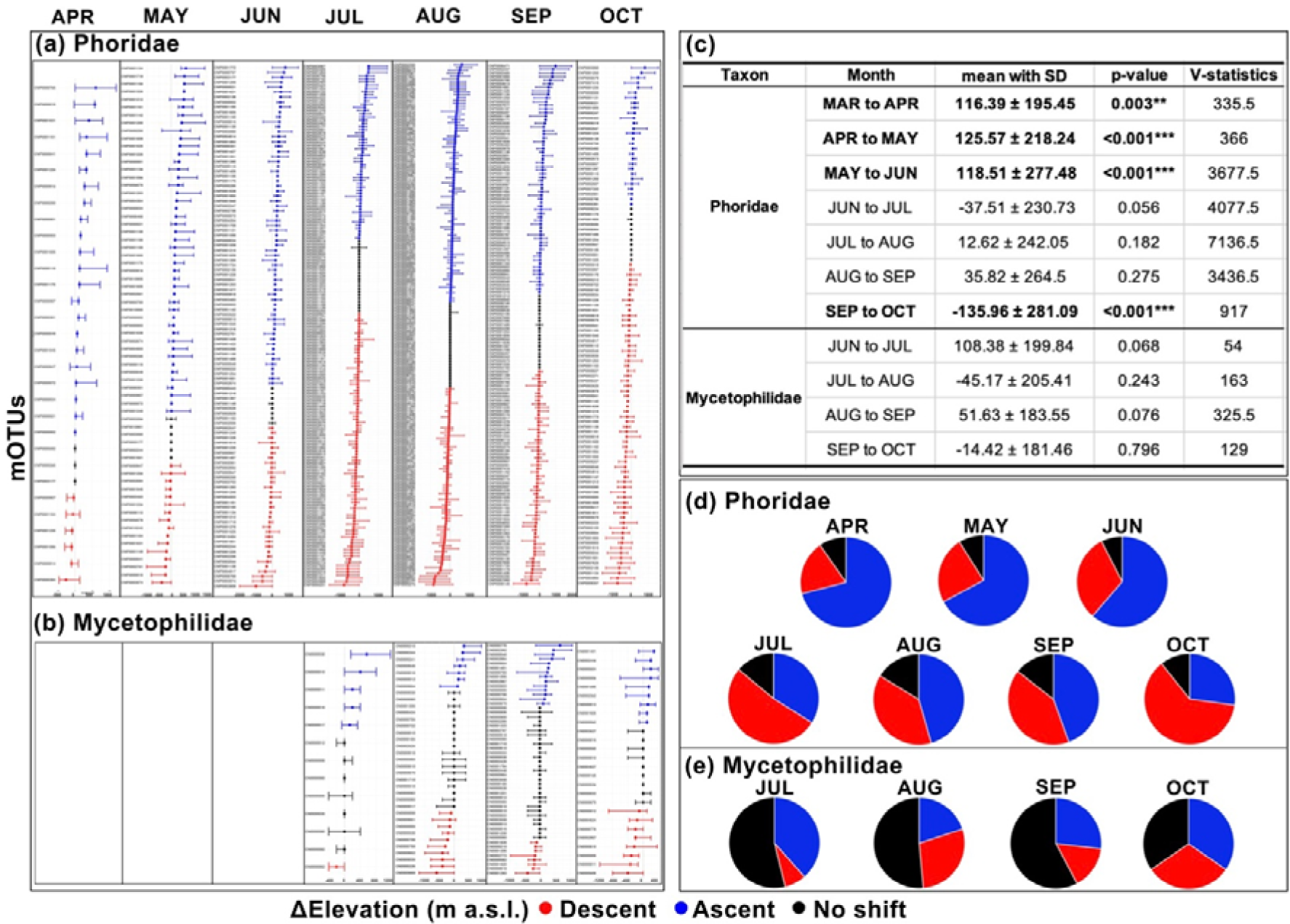
Elevation range shifts for mOTUs with at least one specimen observed in two consecutive months for (a) Phoridae and (b) Mycetophilidae. Months in the figure are in comparison to the previous month. Blue indicates an ascent, red indicates a descent, and black indicates no change in the weighted mean elevation. Dots = weighted mean elevations; whiskers = absolute values of shift of the lowest (left) or highest (right) elevations. (c) Wilcoxon signed-rank tests for significant differences in the weighted mean elevation of consecutive months for Phoridae and Mycetophilidae (p-value: * p < 0.05; ** p < 0.01; *** p < 0.001). Proportion of ascending, descending or unchanged mOTUs for (d) Phoridae and (e) Mycetophilidae.

In addition, the general shifts are also evident in the ranges of individual mOTUs (Figure 5a,b). On average, elevation range shifts between months exceeded 100 m a.s.l. for all significant comparisons (Table S9) and were often even greater for specific mOTUs (Figure 5a,b). Notably, we also found that a higher proportion of mOTUs of Mycetophilidae showed no change in elevation as the months changed (Figure 5b,e). This is very different to what is found for Phoridae (Figure 5a,d).

We subsampled the Phoridae specimens in the month of August which represented the largest number of mOTUs observed across two consecutive months. Random subsampling of Phoridae specimens in August five times resulted in subsets ranging from 1,776 to 3,061 specimens consisting of 35 mOTUs corresponding to the 35 Mycetophilidae mOTUs in August. Random subsampling of Phoridae specimens in August five times resulted in subsets ranging from 133 to 145 mOTUs consisting of 932 specimens corresponding to the 932 Mycetophilidae specimens in August. The same pattern was observed for all phorid subsets (Figures S5 and S6).

### 3.6 Community composition across month and elevation

Phorid communities were strongly grouped according to month while mycetophilid communities were grouped according to the elevation (Figure 6). Figure 6a shows the NMDS plot of phorid communities for all months, and a PERMANOVA test on the NMDS1 & NMDS2 coordinates found significant differences in communities across months (R^2^ = 0.34, p < 0.001) (Table S10). Phorid communities are divided into three distinct temporal periods according to spring, summer and autumn (Figure 6a). The greater the distance between months, the greater the difference in community composition (Table S10). Five phorid subsets ranging from 3,488 to 5,740 specimens consisting of 148 mOTUs and five phorid subsets ranging from 275 to 285 mOTUs consisting of 2,281 specimens were created and the three temporal periods remained clearly distinct (Figures S7 and S8). However, no significant differences between communities were found for Phoridae at different elevations (R^2^ = 0.11, p = 0.397). On the other hand, mycetophilid communities across elevations differed significantly (R^2^ = 0.41, p < 0.001) across months (R^2^ = 0.30, p = 0.002) (Table S10). Communities at middle elevations (1,200 m and 1,400 m a.s.l) in summer months (July and August) showed a tendency to cluster, while communities at low (800 m and 1,000 m a.s.l.) and high (1,800 m a.s.l.) elevations were very distant. Communities found at 1,600 m a.s.l. also tended to cluster (Figure 6b). The above results for Phoridae and Mycetophilidae are robust to the use of trimmed datasets (Figures S9-S12).

**FIGURE 6.**
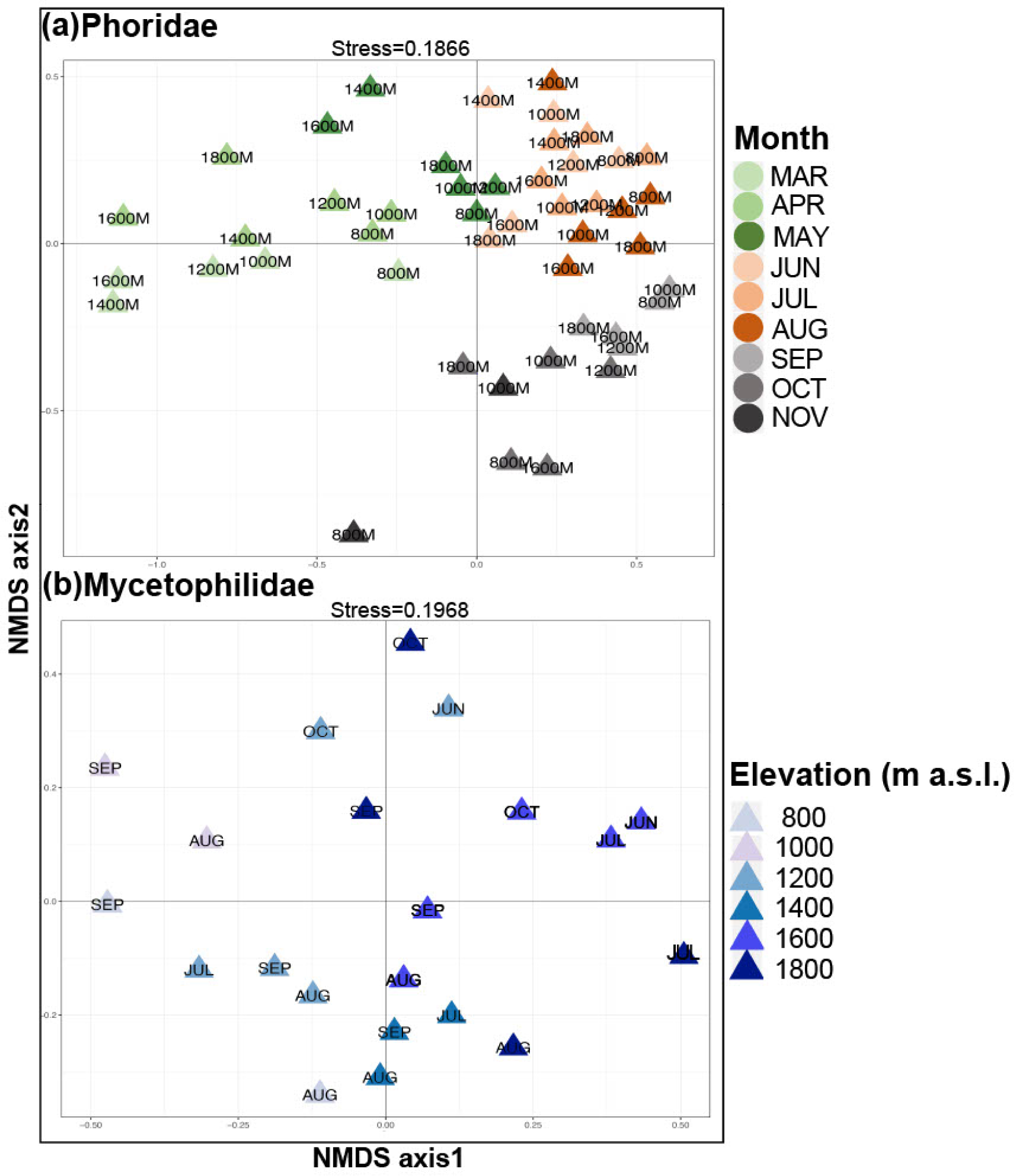
Nonmetric Multidimensional Scaling (NMDS) based on Bray-Curtis dissimilarity matrices illustrated on two-dimensional NMDS plots. This figure uses the full dataset at 4-week intervals, but samples with less than 10 specimens are removed.

For Phoridae, only 92 of the 492 mOTUs (18.70%) occur in all seasons, and 240 mOTUs (48.78%) occur only in a single season (Figure 7a). The largest number of unique mOTUs (174) were found in summer. Furthermore, only seven mOTUs are shared between spring and autumn despite similar temperature and precipitation levels. For Mycetophilidae, only 5 of the 148 mOTUs (3.38%) occur in all elevations while 67 mOTUs (45.27%) occur only in a single elevation (Figure 7b). The largest number of unique mOTUs (20) were found at 1,600 m a.s.l.

**FIGURE 7.**
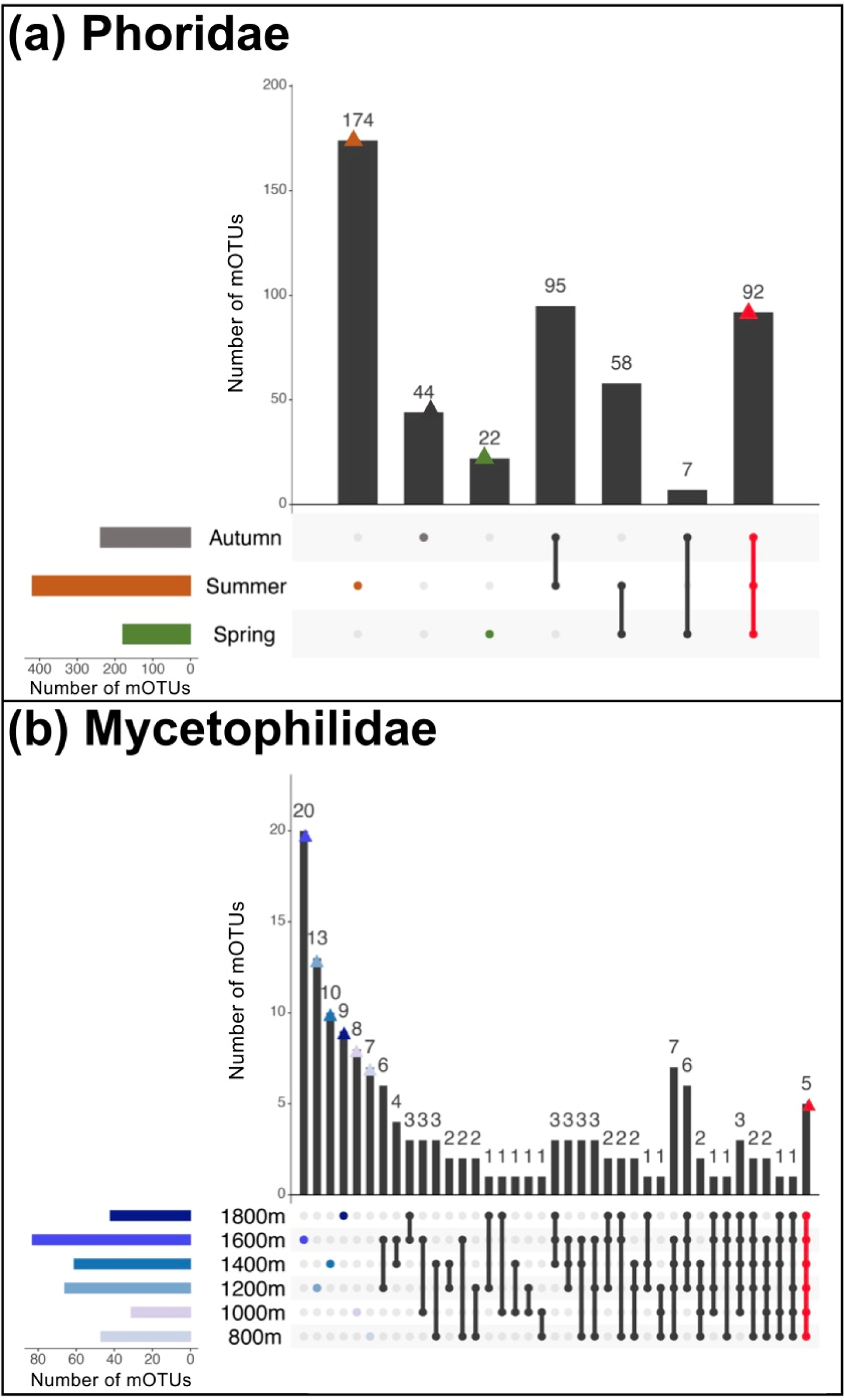
Number of mOTUs unique to and shared across (a) three seasons of Phoridae (Spring: March-May, Summer: June-August, Autumn: September-November) and (b) six elevations of Mycetophilidae.

Phoridae had a higher overall dissimilarity value across months (β_BC_ = 0.871) than Mycetophilidae (β_BC_ = 0.804) (Table 1). However, spatial turnover accounted for most of the dissimilarity in Phoridae (β_BC.BAL_ = 0.781, β_BC.GRA_ = 0.090) while nestedness contributed more in Mycetophilidae (β_BC.BAL_ = 0.372, β_BC.GRA_ = 0.431) (Table 1). In comparison, Mycetophilidae had a higher overall dissimilarity across elevations (β_BC_ = 0.839) than Phoridae (β_BC_ = 0.808) (Table 2). Spatial turnover accounted for most of the dissimilarity in both taxa but to a greater extent in Mycetophilidae (β_BC.BAL_ = 0.721, β_BC.GRA_ = 0.118, β_BC.BAL/_β_BC_ ratio = 0.859) than in Phoridae (β_BC.BAL_ = 0.683, β_BC.GRA_ = 0.125, β_BC.BAL/_β_BC_ ratio = 0.845) (Table 2). These results are robust to removal of rare mOTUs (Table S11).

**Table 1.**
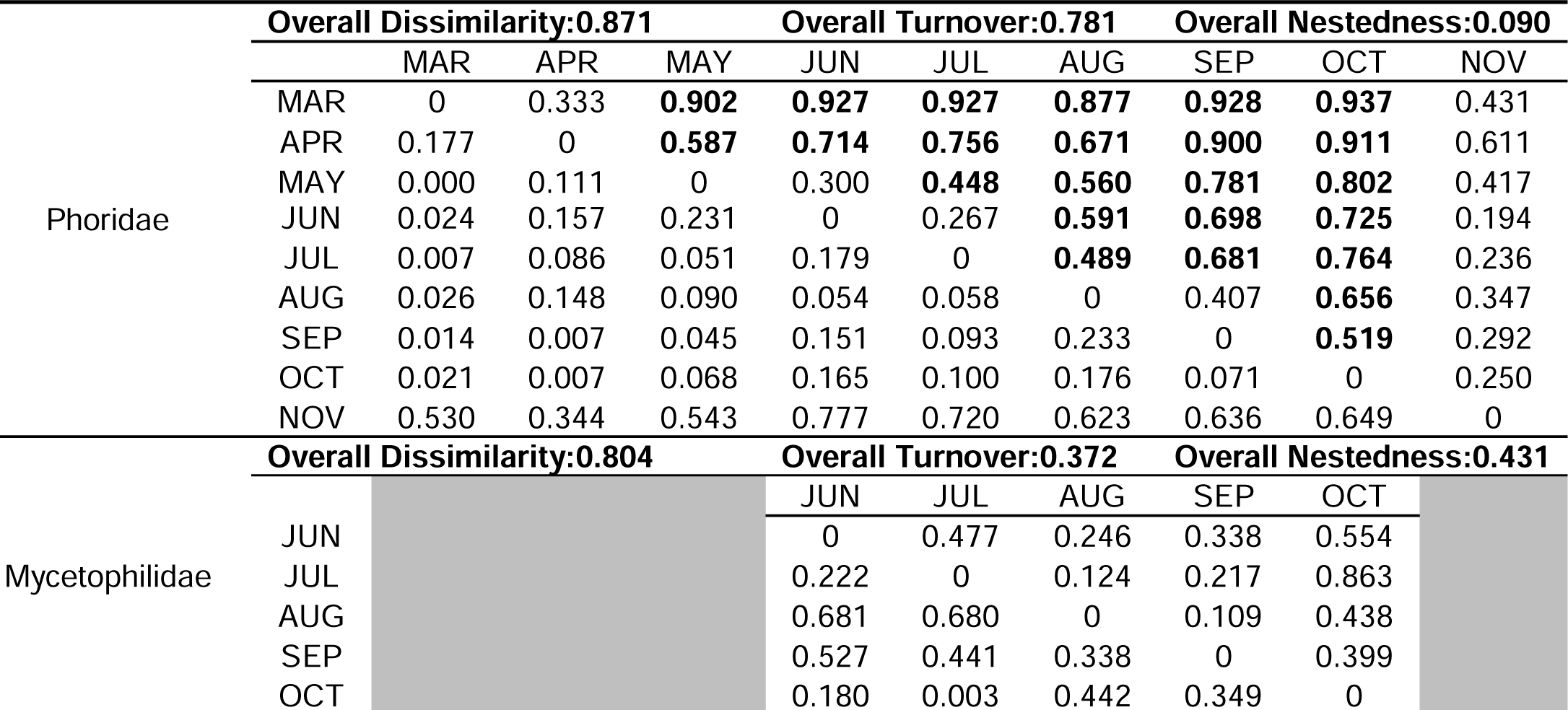
Beta diversity of Phoridae and Mycetophilidae across months. Multiple-site dissimilarity values and their partitions (turnover and nestedness) are indicated at the top of each table. Pairwise turnover values are above and pairwise nestedness values are below the diagonal. Bold numbers indicate that the turnover is three or more times that of the nestedness. Calculations are based on the full datasets at 4-week intervals, with removal of samples with less than 10 specimens.

**Table 2.** Beta diversity of Phoridae and Mycetophilidae across elevations. Multiple-site dissimilarity values and their partitions (turnover and nestedness) are indicated at the top of each table. Pairwise turnover values are above and pairwise nestedness values are below the diagonal. Bold numbers indicate that the turnover is three or more times that of the nestedness. Calculations are based on the full datasets at 4-week intervals, with removal of samples with less than 10 specimens.

|  |  | Overall Dissimilarity:0.808 |  | Overall Turnover:0.683 |  | Overall Nestedness:0.125 |  |
| --- | --- | --- | --- | --- | --- | --- | --- |
|  |  | 800M | 1000M | 1200M | 1400M | 1600M | 1800M |
| Phoridae | 800M | 0 | 0.208 | **0.528** | 0.330 | **0.615** | **0.746** |
|  | 1000M | 0.323 | 0 | **0.576** | 0.220 | **0.716** | **0.579** |
|  | 1200M | 0.076 | 0.112 | 0 | 0.199 | **0.605** | **0.643** |
|  | 1400M | 0.346 | 0.595 | 0.500 | 0 | 0.356 | **0.611** |
|  | 1600M | 0.093 | 0.052 | 0.033 | 0.434 | 0 | **0.426** |
|  | 1800M | 0.017 | 0.194 | 0.080 | 0.181 | 0.174 | 0 |
|  |  | Overall Dissimilarity:0.839 |  | Overall Turnover:0.721 |  | Overall Nestedness:0.118 |  |
|  |  | 800M | 1000M | 1200M | 1400M | 1600M | 1800M |
| Mycetophilidae | 800M | 0 | **0.471** | 0.256 | 0.519 | 0.431 | **0.663** |
|  | 1000M | 0.073 | 0 | 0.438 | **0.835** | **0.669** | **0.636** |
|  | 1200M | 0.421 | 0.367 | 0 | **0.652** | **0.590** | **0.519** |
|  | 1400M | 0.267 | 0.106 | 0.006 | 0 | **0.711** | **0.707** |
|  | 1600M | 0.340 | 0.225 | 0.020 | 0.019 | 0 | 0.448 |
|  | 1800M | 0.021 | 0.072 | 0.251 | 0.149 | 0.308 | 0 |

### 3.7 Mid-domain effect

Model predictions of MDE model for both Mycetophilidae and Phoridae trended towards a mid-domain peak, as expected (Figure S13). However, Phoridae exhibited a sharp decline in richness at 1,400 m a.s.l., with a primary richness peaking at 1,200 m a.s.l. and a smaller secondary peak at 1,600 m a.s.l. In contrast, Mycetophilidae displayed a singular peak in richness at 1,600 m a.s.l.

### 3.8 Phylogenetic structure

Based on the NTI obtained, Phoridae showed slight phylogenetic clustering at low and middle elevation (800 m and 1,400 m a.s.l.: NTI > 2) and strong clustering at high elevations (1,600 m and 1,800 m a.s.l.: NTI > 4). Mycetophilidae showed a phylogenetic clustering at the highest elevation (1,800 m a.s.l.: NTI > 2) (Table S12a). For seasonal communities, Phoridae showed phylogenetic clustering in spring (March, April and May: NTI > 2), part of summer (August: NTI > 2) and autumn (September, October and November: NTI > 2) whereas Mycetophilidae only showed strong clustering in part of summer (June: NTI > 2) (Table S12b). There was no overdispersion in both taxa along elevation and month community. Note that these results are only based on DNA Barcode data so that relationships and branch lengths could only be approximated.

## DISCUSSION

### Very different tales for two dark taxa

Little is known about the spatio-temporal distribution of most animal species that are concentrated in comparatively poorly known dark taxa. In particular, those species belonging to dark taxa are so poorly known that they are species-rich versions of the more charismatic faunas such as the ones of the deep sea, groundwater, or canopy. Yet, these dark taxa live all around us and we should be interested in having a sounder understanding of their ecosystem services. Sampling is not that the problem because they regularly comprise more than 50% of the specimens in Malaise trap samples (Srivathsan et al., 2023). The problem is the lack of analysis at the specimen level. We here demonstrate that this problem can be overcome with megabarcoding which not only yields alpha and beta diversity data, but also abundance information that allows for the use of a broader range of analysis tools. Once applied, we find that the two dark taxa covered here, Mycetophilidae and Phoridae, have quite different spatio-temporal distributions despite having been sampled by the same traps at the same sites.

Firstly, the diversity and abundance of Phoridae is vastly higher than that of Mycetophilidae (richness: 492 vs 148; abundance: 17,179 vs 2,281). This was somewhat expected as Srivathsan et al., 2023 found Phoridae to be three times more abundant than Mycetophilidae in Malaise traps globally. Our results indicate that the relative difference in abundance can be even larger, but the two taxa have even clearer differences in richness and abundance over time (Figures 2 and 3). Phorids appear earlier and disappear later in the seasons. This may be due to the great ecological diversity of the taxon (Disney, 1994; Srivathsan et al., 2019). Many species have saprophagous larvae, but others are herbivores, fungivores, predators, or even parasites/parasitoids (Disney, 1979, 1990, 1994; Kitching et al., 2005; Ronquist et al., 2020), sometimes with very specific adaptations (Weissflog et al., 1995). The family is also known to play a significant role in colonizing habitats (Weber & Prescher, 1990; Durska, 2001, 2006; Prescher et al., 2002) suggesting that they can thrive under a great variety of ecological conditions. Other species are adapted to extremely cold climates, thriving well above the Arctic Circle (Disney, 2004), or enduring winter conditions in montane areas (Mostovski & Disney, 2002). This may explain why the phorids of the Baihua Mountain Reserve disappear later in the year than the mycetophilids. We furthermore find evidence for strong phylogenetic clustering at high elevations, i.e., the potential for a high-elevation clade of species although phylogenetic conclusions drawn from COI data only should be interpreted with care.

On the contrary, Mycetophilidae were only observed one month after the spring rains, and they were generally not observed in low-precipitation months. Instead, alpha diversity and abundance increased gradually with increasing precipitation, with a distinct temporal lag between precipitation and richness peaks (Figure 4). The feeding habits of adult mycetophilids are poorly known, but the larvae for most species are linked to fungi, hyphae, or decaying wood (Ševčík, 2010; Jakovlev, 2012; Kaspřák et al., 2018). Precipitation is closely related to vegetation and fungal growth (Zak et al., 2003; Hawkes et al., 2011), thus providing more habitat and food resources for mycetophilids. A period of insect reproduction and development may lead to peak adult insect communities occurring in the weeks or months following precipitation (Dietrich et al., 2023). Host species diversity have been recognized as crucial factors influencing consumer richness, particularly among herbivores and their parasitoids (Elizalde & Folgarait, 2010). Mushrooms, in particular, are an ephemeral and unpredictable food sources, supporting polyphagous feeders (Hanski, 1989). Unfortunately, empirical studies investigating the relationships within the fungi-fungivore community and the differentiation between specialist and generalist fungivores remain scarce (Põldmaa et al., 2015) and there is thus limited information on how fungal diversity may influence local fungus gnat communities. Our detailed information on the spatio-temporal distribution of Mycetophilidae of the Baihua Mountain Reserve will help with designing a complementary study of fungal diversity to understand these interdependencies. Such a follow-up study is warranted because the mycetophilid community of Baihua Mountain Reserve appears dependent on the interplay between precipitation and temperature patterns, i.e., environmental factors that are likely changing rapidly as we report these results.

We observed a significant turnover in mOTUs for the two dark taxa across different months and elevations, suggesting a specialization of these communities for specific time periods and elevations. However, these compositional changes in community structure differ between the two dark taxa. Seasonality is important for phorid communities, while mycetophilid communities were uniform across months, possibly due to their overall time of activity also being markedly shorter (Figures 5 and 6, Tables S10 and S11). Instead, mycetophilid communities differed more along the elevational gradient than phorid communities.

Phorids showed three distinct communities, one in spring (March to May), summer (June to August) and autumn (September to November) (Figure 6) with gradual turnover from one month to the next. In contrast, very little structure was caused by elevation. This suggests that the community as a whole shifts potentially due to relatively large ranges and no strict attachment to a habitat, but they may be more dependent on food resources, which may be more widespread. This is consistent with our observation that phorids generally had a smaller proportion of mOTUs found at only one elevation and a larger proportion of mOTUs to shift their distribution to higher or lower elevation dependent on season. Generally, an upwards movement of mOTUs was observed from spring to summer which was replaced by a downward movement from summer to autumn (Figure 5). Interestingly, despite similar temperatures and levels of precipitation in the spring and autumn seasons, a very low number of mOTUs are shared between these two seasons (Figure 7a).

Mycetophilids are dependent on more ephemeral resources in the form of fungi, and their community composition is more influenced by elevation (Figure 6). This could be due to being very specialized on specific fungus that may be clumped (Põldmaa et al., 2015; Kaspřák et al., 2018) at specific elevations. This is supported by our results that show a much smaller proportion of mycetophilid mOTUs found at all elevations (Figure 7b). Indeed, our results suggest based on our very limited data that mycetophilid species in Baihua Mountain Reserve could have small ranges, with very few mOTUs being found at different elevations across months (Figure 5). Mycetophilids are known to be very sensitive to moisture and shade and probably only few specialized species are able to inhabit the disturbed low-elevation areas and subalpine meadows both with higher light intensity, which compared to the mid-elevation areas with well-developed lower canopy showed lower diversity of mycetophilids.

Most studies of the mid-domain effect (MDE) use well-known taxa, for which elevational distributions have been established over decades, e.g. plants or vertebrates (Jetz & Rahbek, 2002; Colwell et al., 2016; Liu et al., 2019; He et al., 2019). Comparatively little is known about most insect groups including dark taxa although the latter are so abundant and diverse that they could be a rich data source. We here study the MDE based for phorids or mycetophilids, that are likely to have a fairly low dispersal ability (Jakovlev, 2012; Disney, 1994) compared to many vertebrate groups but are more likely to respond to change faster than most plant species. For both taxa we find some evidence for a mid-domain distribution pattern (Figure S13), although the data are not conclusive. Note that such conflicting support (Colwell et al., 2004) can be due to veil effects resulting from incomplete sampling with only six elevations sampled for one year. Also, not sampling enough sites at each elevation, or not sampling at enough elevations will result in imprecisely estimated species ranges. This is a common challenge to studying species richness patterns along elevational gradients in terrestrial ecosystems (Nogués-Bravo et al., 2008). That said this provides first observation of MDE patterns in two highly diverse and understudied groups. More intense sampling across multiple years can help determine if the observed trend become clearer and if similar observation can be made for other groups.

### Drivers of the elevation shifts

Monitoring of spatio-temporal distribution patterns of phorids and mycetophilids at different elevations could provide important information on climatic changes in the reserve. We observed that phorid species shift along elevation gradients in the Baihua Mountain Reserve. Many mOTUs appear earlier in the annual cycle at lower elevations and later at higher elevations (Figure 3, Table S2). For example, we observed that mOTU CNP0001161 was first collected in 21-28 March at 800 m a.s.l, with more specimens occurring at higher elevations over time (Tables S2 and S4). The phenology patterns were different at different elevations (e.g. mOTU CNP0001235, Table S2). The differences are likely due to temperature and precipitation. We hypothesize that the seasonal shifts in both richness and elevation ranges could be due to the delayed emergence of adults at higher elevations and/or the seasonal migration of individuals. Phenological variations in mountain insects are primarily influenced by seasonal temperature changes (Bishop et al., 2014; Boulter, Lambkin, & Starick, 2011; Wardhaugh, Stone, & Stork, 2018). This is because the lower temperatures of higher elevations affect the metabolism and life activities of insects, causing them to move more slowly and undergo a prolonged development period, ultimately resulting in the later emergence of adults. Temperature and precipitation also alter the availability of food resources, which in turn can trigger seasonal migrations of adults (Janzen, 1987; Hsiung et al., 2018; Maicher et al., 2020). Depletion of these resources in one area may prompt insects to migrate elsewhere in search of more abundant food resources.

Detailed studies of temporal change of diversity along elevation gradients are scarce because of the difficulty of long-term and continuous sampling at all elevations. Inconsistent results are common, whether between different studies or within the same taxa (Janzen,1973; Wardhaugh et al., 2018; Hsiung et al., 2018; Maicher et al., 2018, 2020; Beck & Linsenmair, 2006; Owen, 2009). For instance, Janzen (1973) demonstrated a decline in beetle diversity at mid-elevations during the wet season. However, a study in the wet tropics of Australia by Wardhaugh et al. (2018) revealed no consistent seasonal shifts in beetle abundance or richness along elevation. The study by Maicher et al. (2020) on tropical Lepidoptera found that most species displayed an upward shift from beginning to end of the dry season, but Arctiinae shifted upwards during the wet season. Of course, comparable data for phorids and mycetophilids are completely missing but our results highlight the importance of considering temporal dynamics when interpreting elevation patterns of diversity in the context of climate change. Understanding the changes of species richness in different global ecosystems over time will be needed for providing guidance for future biodiversity conservation (White et al., 2006; Suurkuukka et al., 2012).

### Revealing detailed insect biodiversity patterns with megabarcoding

The ability to establish spatio-temporal distribution patterns for large numbers of neglected insect species at once can be a major step toward understanding insect biodiversity in greater depth. High-throughput specimen-based DNA barcoding could contribute significantly to addressing the DNA barcode reference data gap for dark taxa, accelerating species discovery (Srivathsan et al., 2023, Fernandez-Triana, 2022), generating detailed insect biodiversity patterns (Baloğlu et al., 2018; Yeo et al., 2021; Geiger et al., 2016), allowing for rapid biodiversity species inventories (Telfer et al., 2015), and tracking baseline shifts (D’Souza et al., 2021). The benefits include a link between specimen and barcode, giving insights into abundances and phylogenetic relationships of the studied communities. We show that accurate data on abundance and distribution may be obtained for habitat assessments using NGS barcodes (Meier et al., 2016, Wang et al., 2018; Yeo et al., 2021; Hartop et al., 2022; D’souza & Hebert, 2018). Using efficient and cost-effective techniques, one lab member can amplify the barcodes for 600-1,000 specimens per day. Species discovery can be rapid. For example, in this study we found 492 putative phorid and 148 mycetophilid species, while the number of known species in China is only 226 and 260 species of phorids and mycetophilids respectively (Liu et al., 2023; Ye et al., 2023). This means that studying only one elevational gradient in a temperate reserve for phorids more than doubles the number of putative species known for a very large and biodiverse country. Implications for studies of global change

Montane ecosystems are some of the most threatened on the planet (La Sorte & Jetz 2010; Goodenough & Hart, 2013; Sheldon et al., 2011; De Gabriel Hernando et al., 2021). The current consensus is that climate change will cause upwards shifts of tropical montane species (Forero-Medina et al., 2011; Laurance et al., 2011; Chen et al., 2009), resulting in subsequent extinctions of mountaintop species (Colwell et al., 2008; Freeman et al., 2021). For the biodiversity of temperate mountain forests, species adapted to cold-climatic mountain environments are expected to face a high risk of range contractions ultimately causing local extinctions (Hughes, 2000; Thomas et al., 2004; Braunisch et al., 2014). We observed some phylogenetic clustering for phorid and mycetophilid species in high elevation communities, thus implying a potential threat to the survival of these taxa. However, our study also reveals that many mOTUs shift more than 100 m, with some mOTUs having changes several times higher. This suggests that many other species could be quite tolerant to climatic changes.

## CONCLUSION

Analysis of biodiversity patterns of dark taxa through acceleration of species discovery is an urgent task, because climate change is likely threatening the very large number of species in these taxa. Our results indicate the importance of conducting detailed spatio-temporal studies of diversity distribution patterns for multiple groups. The two dark taxa covered here follow very different patterns, but both were impressively rich and distinct along the temporal, spatial, and elevational gradient. Our finding indicates that megabarcoding is a technically and economically feasible approach to answering ecological questions granting us more detailed insights into biodiversity patterns of unknown, oft-neglected megadiverse groups. Future studies should consider including classical approaches in combination with megabarcoding techniques to allow further insights into the differences between dark taxa and diversity distribution patterns of different taxa.

## AUTHOR CONTRIBUTIONS

W.P., V.F., A.H., A.S., D.Z. and R.M. conceived and designed the research study. W.P., L.Y., W.X., H.S., J.Y., X.Z., N.Y. and Q.Y. collected the samples. W.P. generated and analyzed the data with guidance from V.F., A.K.H., A.S., D.Z. and R.M. W.P., V.F., A.K.H., D.Z. and R.M. wrote the manuscript with revisions from all authors.

## ACKNOWLEDGMENTS

This work was supported by the Center for Integrative Biodiversity Discovery, Leibniz Institute for Evolution and Biodiversity Science, Museum für Naturkunde Berlin (Rudolf Meier). W.P. acknowledges support from China Scholarship Council for her study at the Museum für Naturkunde Berlin (No. 202006510035). A.K.H. acknowledges the Carlsberg Foundation for their continuous support of his postdoc activities through the project ‘Next Generation Taxonomy’. Special thanks to Prof. Dr. Dalton de Souza Amorim and Dr. Daniel Dias Dornelas do Carmo of the Universidade de São Paulo and Prof. Dr. Diego Aguilar Fachin of the Universidade Federal de Goiás for verifying the specimens of Mycetophilidae in this paper.

## DATA AVAILABILITY STATEMENT

The data are available from https://figshare.com/account/home#/projects/197716

