## Supplementary Tables S6-S10 for "Megabarcoding reveals a tale of two very different dark taxa along the same elevational gradient"

**Supplemental Table S6.** Number and distribution of mOTUs delimited using different thresholds (19,460 barcoded specimens).

| **Taxon** | **No. of Barcodes** | **No. of mOTUs from Objective Clustering** | | |
| --- | --- | --- | --- | --- |
|  |  | **2%** | **3%** | **4%** |
| **Phoridae** | 17,179 | 537 | 492 | 451 |
| **Mycetophilidae** | 2,281 | 151 | 148 | 144 |
| **Total** | 19,460 | 688 | 640 | 595 |

**Supplemental Table S7.** Diversity of Phoridae in different elevations and months in the Baihua Mountain Reserve.

| Month | Elevation  (m a.s.l.) | Richness | Abundance | Sample coverage |
| --- | --- | --- | --- | --- |
| MARCH | 800 | **51** | 616 | 0.96 |
|  | 1000 | 23 | **1020** | **0.99** |
|  | 1200 | 13 | 214 | 0.97 |
|  | 1400 | 5 | 116 | 1.00 |
|  | 1600 | 6 | 46 | 0.96 |
|  | 1800 | 3 | 3 | 0.33 |
| APRIL | 800 | 36 | 253 | 0.95 |
|  | 1000 | **45** | **483** | 0.97 |
|  | 1200 | 30 | 189 | 0.93 |
|  | 1400 | 17 | 161 | 0.92 |
|  | 1600 | 5 | 33 | **0.97** |
|  | 1800 | 13 | 49 | 0.88 |
| MAY | 800 | 60 | 285 | 0.90 |
|  | 1000 | **83** | **825** | **0.96** |
|  | 1200 | 87 | 368 | 0.92 |
|  | 1400 | 24 | 40 | 0.58 |
|  | 1600 | 23 | 88 | 0.89 |
|  | 1800 | 72 | 426 | 0.93 |
| JUNE | 800 | 75 | 434 | 0.92 |
|  | 1000 | 65 | 1028 | **0.97** |
|  | 1200 | 102 | 765 | 0.95 |
|  | 1400 | 39 | 111 | 0.79 |
|  | 1600 | **112** | **1233** | 0.97 |
|  | 1800 | 77 | 453 | 0.93 |
| JULY | 800 | 66 | 233 | 0.82 |
|  | 1000 | 94 | **765** | **0.94** |
|  | 1200 | **111** | 385 | 0.88 |
|  | 1400 | 69 | 174 | 0.75 |
|  | 1600 | 99 | 516 | 0.92 |
|  | 1800 | 81 | 376 | 0.90 |
| AUGUST | 800 | 77 | 266 | 0.83 |
|  | 1000 | 94 | 759 | 0.95 |
|  | 1200 | 139 | 435 | 0.86 |
|  | 1400 | 38 | 112 | 0.78 |
|  | 1600 | **148** | **1202** | **0.95** |
|  | 1800 | 60 | 305 | 0.92 |
| SEPTEMBER | 800 | 48 | 94 | 0.69 |
|  | 1000 | 59 | 222 | 0.86 |
|  | 1200 | 75 | 231 | 0.82 |
|  | 1400 | 5 | 6 | 0.39 |
|  | 1600 | **106** | **522** | **0.92** |
|  | 1800 | 64 | 267 | 0.89 |
| OCTOBER | 800 | 35 | 63 | 0.59 |
|  | 1000 | 57 | 235 | 0.90 |
|  | 1200 | **70** | **539** | **0.93** |
|  | 1400 | 2 | 2 | 0.67 |
|  | 1600 | 26 | 65 | 0.75 |
|  | 1800 | 42 | 94 | 0.72 |
| NOVEMBER | 800 | 13 | 20 | 0.57 |
|  | 1000 | 25 | 44 | 0.59 |
|  | 1200 | 1 | 1 | 1.00 |
|  | 1400 | 0 | 0 | - |
|  | 1600 | 0 | 0 | - |
|  | 1800 | 6 | 7 | 0.33 |

Note. The highest values of each diversity measure for each month are indicated in bold.

**Supplemental Table S8.** Diversity of Mycetophilids in different elevations and months in the Baihua Mountain Reserve.

| Month | Elevation  (m a.s.l.) | Richness | Abundance | Sample coverage |
| --- | --- | --- | --- | --- |
| MARCH | 800 | 0 | 0 | - |
|  | 1000 | 0 | 0 | - |
|  | 1200 | 0 | 0 | - |
|  | 1400 | 0 | 0 | - |
|  | 1600 | 0 | 0 | - |
|  | 1800 | 0 | 0 | - |
| APRIL | 800 | 0 | 0 | - |
|  | 1000 | 0 | 0 | - |
|  | 1200 | 0 | 0 | - |
|  | 1400 | 0 | 0 | - |
|  | 1600 | 0 | 0 | - |
|  | 1800 | 0 | 0 | - |
| MAY | 800 | 1 | 1 | 1 |
|  | 1000 | 0 | 0 | - |
|  | 1200 | 1 | 1 | 1 |
|  | 1400 | 0 | 0 | - |
|  | 1600 | 0 | 0 | - |
|  | 1800 | 0 | 0 | - |
| JUNE | 800 | 2 | 3 | 0.83 |
|  | 1000 | 1 | 1 | 1 |
|  | 1200 | **14** | **42** | **0.86** |
|  | 1400 | 3 | 4 | 0.63 |
|  | 1600 | 10 | 23 | 0.75 |
|  | 1800 | 5 | 8 | 0.66 |
| JULY | 800 | 3 | 5 | 0.73 |
|  | 1000 | 5 | 9 | 0.59 |
|  | 1200 | 12 | 44 | 0.82 |
|  | 1400 | **25** | **86** | **0.83** |
|  | 1600 | 10 | 16 | 0.58 |
|  | 1800 | 6 | 15 | 0.88 |
| AUGUST | 800 | 39 | 120 | 0.83 |
|  | 1000 | 23 | 101 | 0.86 |
|  | 1200 | 45 | **350** | 0.94 |
|  | 1400 | 37 | 323 | **0.95** |
|  | 1600 | **62** | 313 | 0.92 |
|  | 1800 | 25 | 70 | 0.79 |
| SEPTEMBER | 800 | 9 | 40 | 0.88 |
|  | 1000 | 7 | 20 | 0.76 |
|  | 1200 | 30 | 128 | 0.86 |
|  | 1400 | 27 | 149 | **0.93** |
|  | 1600 | **36** | **164** | 0.89 |
|  | 1800 | 20 | 75 | 0.88 |
| OCTOBER | 800 | 4 | 4 | 0.18 |
|  | 1000 | 6 | 7 | 0.33 |
|  | 1200 | 8 | 13 | 0.63 |
|  | 1400 | 4 | 5 | 0.49 |
|  | 1600 | **17** | **120** | **0.93** |
|  | 1800 | 9 | 20 | 0.82 |
| NOVEMBER | 800 | 1 | 1 | 1 |
|  | 1000 | 0 | 0 | - |
|  | 1200 | 0 | 0 | - |
|  | 1400 | 0 | 0 | - |
|  | 1600 | 0 | 0 | - |
|  | 1800 | 0 | 0 | - |

Note. The highest values of each diversity measure for each month are indicated in bold.

**Supplemental Table S9.** Detailed results of the Wilcoxon signed-rank tests of the changes in the three measures of elevational range (highest elevation, weighted mean elevation and lowest elevation) for phorids and mycetophilids. The first number gives the V-statistic, while the second number represents the p-value (* p < 0.05; ** p < 0.01; *** p < 0.001), and the third number represents the mean change in elevation with standard deviations given.

**All mOTUs with at least one specimen in two consecutive months**

| Range measure | Month | Values | Phoridae | Mycetophilidae |
| --- | --- | --- | --- | --- |
| Highest elevation | March to April | V; p-value; mean with SD | 152.5, 0.003**, 300 ± 316.23 | - |
|  | April to May |  | 20.5, <0.001***, 334.33 ± 352.29 | - |
|  | May to June |  | 973, 0.359, 20.75 ± 341.35 | - |
|  | June to July |  | 1509, 0.471, -15.85 ± 324.83 | 38，0.066, 138.46 ± 236.43 |
|  | July to August |  | 2186, 0.112, 24.21 ± 293.21 | 136, 0.097, 80 ± 262.12 |
|  | August to September |  | 1505, 0.189, -30.34 ± 331.1 | 173.5, 0.494, -19.23 ± 202.96 |
|  | September to October |  | 370.5, <0.001***, -132.04 ± 347.04 | 20, 0.07, -76.92 ± 204.56 |
| Weighted mean elevation | March to April | V; p-value; mean with SD | 335.5, 0.003**, 116.39 ± 195.45 | - |
|  | April to May |  | 366, <0.001***, 125.57 ± 218.24 | - |
|  | May to June |  | 3677.5, <0.001***, 118.51 ± 277.48 | - |
|  | June to July |  | 4077.5, 0.056, -37.51 ± 230.73 | 54，0.068, 108.38 ± 199.84 |
|  | July to August |  | 7136.5, 0.182, 12.62 ± 242.05 | 163, 0.243, -45.17 ± 205.41 |
|  | August to September |  | 3436.5, 0.275, 35.82 ± 264.5 | 325.5, 0.076, 51.63 ± 183.55 |
|  | September to October |  | 917, <0.001***, -135.96 ± 281.09 | 129, 0.796, -14.42 ± 181.46 |
| Lowest elevation | March to April | V; p-value; mean with SD | 48, 0.484, 32.26 ± 200.64 | - |
|  | April to May |  | 322.5, 0.135, -44.78 ± 257.78 | - |
|  | May to June |  | 1248, <0.001***, 116.98 ± 342.39 | - |
|  | June to July |  | 1719, 0.033*, -41.46 ± 272.66 | 31，0.753, 30.77 ± 268.9 |
|  | July to August |  | 2805, 0.806, 3.16 ± 299.9 | 12，<0.001***, -240 ± 264.8 |
|  | August to September |  | 1232.5, 0.013*, 85.52 ± 387.83 | 108, 0.006**, 130.77 ± 331.09 |
|  | September to October |  | 811.5, 0.122, -58.25 ± 364.75 | 132, 0.316, 69.23 ± 391.68 |

**No Doubletons**

| Range measures | Month | Value | Phoridae | Mycetophilidae |
| --- | --- | --- | --- | --- |
| Highest elevation | March to April | V; p-value; mean with SD | 123, 0.004**, 160 ± 274.93 | - |
|  | April to May |  | 20.5, <0.001***, 339.39 ± 352.52 | - |
|  | May to June |  | 948, 0.330, 22.86 ± 342.29 | - |
|  | June to July |  | 1463.5, 0.847, -2.53 ± 317.62 | 33，0.037*, 166.67 ± 222.93 |
|  | July to August |  | 2074, 0.081, 28.11 ± 289.0 | 135, 0.029*, 106.25 ± 243.55 |
|  | August to September |  | 1438, 0.248, -25.53 ± 329.37 | 150.5, 0.423, -20.83 ± 194.56 |
|  | September to October |  | 351, <0.001***, -136.63 ± 348.63 | 20, 0.069, -76.92 ± 204.56 |
| Weighted mean elevation | March to April | V; p-value; mean with SD | 295.5, 0.002**, 100.27 ± 176.59 | - |
|  | April to May |  | 366, <0.001***, 127.47 ± 219.36 | - |
|  | May to June |  | 3638, <0.001***, 121.55 ± 277.04 | - |
|  | June to July |  | 4056, 0.150, -25.01 ± 217.24 | 51，0.019*, 134.08 ± 184.93 |
|  | July to August |  | 6952, 0.143, 16.2 ± 235.62 | 163, 0.368, -30.66 ± 189.73 |
|  | August to September |  | 3315.5, 0.210, 42.51 ± 258.64 | 286, 0.062, 55.94 ± 171.77 |
|  | September to October |  | 865, <0.001***, -140.63 ± 281.52 | 129, 0.796, -14.42 ± 181.46 |
| Lowest elevation | March to April | V; p-value; mean with SD | 36, 0.813, 13.33 ± 173.67 | - |
|  | April to May |  | 322.5, 0.135, -45.45 ± 259.69 | - |
|  | May to June |  | 1214, <0.001***, 120 ± 342.61 | - |
|  | June to July |  | 1685.5, 0.098, -29.11 ± 263.08 | 27，0.626, 50 ± 271.36 |
|  | July to August |  | 2677.5, 0.726, 6.49 ± 296.28 | 12，<0.001***, -243.75 ± 262.66 |
|  | August to September |  | 1157.5, 0.007**, 93.62 ± 385.86 | 86.5, 0.004**, 141.67 ± 332.52 |
|  | September to October |  | 776.5, 0.109, -61.39 ± 367.42 | 132, 0.316, 69.23 ± 391.68 |

**No mOTU <5 Specimens**

| Range measures | Month | Value | Phoridae | Mycetophilidae |
| --- | --- | --- | --- | --- |
| Highest elevation | March to April | V; p-value; mean with SD | 116.5, 0.001**, 185.71 ± 260.65 | - |
|  | April to May |  | 14, <0.001***, 347.69 ± 348.71 | - |
|  | May to June |  | 887, 0.323, 23.76 ± 347.89 | - |
|  | June to July |  | 1347, 0.827, -2.63 ± 321.41 | 33，0.037*, 166.67 ± 222.93 |
|  | July to August |  | 1905.5, 0.066, 32.37 ± 294.71 | 109.5, 0.005**, 150 ± 221.94 |
|  | August to September |  | 1324, 0.158, -34.59 ± 315.76 | 150.5, 0.423, -23.81 ± 208.14 |
|  | September to October |  | 340, <0.001***, -134.02 ± 353.23 | 20, 0.069, -83.33 ± 211.96 |
| Weighted mean elevation | March to April | V; p-value; mean with SD | 264, 0.001**, 113.39 ± 168.31 | - |
|  | April to May |  | 339.5, <0.001***, 130.97 ± 219.2 | - |
|  | May to June |  | 3507.5, <0.001***, 125.37 ± 279.56 | - |
|  | June to July |  | 3894, 0.163, -25.34 ± 217.16 | 51，0.019*, 134.08 ± 184.93 |
|  | July to August |  | 6653.5, 0.090, 20.8 ± 240.23 | 159.5, 0.946, 0.68 ± 170.62 |
|  | August to September |  | 3077, 0.245, 37.8 ± 238.93 | 257, 0.064, 61.55 ± 179.96 |
|  | September to October |  | 845.5, <0.001***, -138.18 ± 285.61 | 113.5, 0.685, -19.79 ± 187.57 |
| Lowest elevation | March to April | V; p-value; mean with SD | 25, 0.803, 14.29 ± 171.52 | - |
|  | April to May |  | 322.5, 0.135, -46.15 ± 261.65 | - |
|  | May to June |  | 1131, <0.001***, 122.77 ± 346.38 | - |
|  | June to July |  | 1577.5, 0.104, -28.95 ± 263.86 | 27，0.626, 50 ± 271.36 |
|  | July to August |  | 2536, 0.560, 11.56 ± 302.48 | 10.5，<0.001***, -235.71 ± 266.96 |
|  | August to September |  | 1010.5, 0.008**, 93.23 ± 380.23 | 73.5, 0.005**, 157.14 ± 346.51 |
|  | September to October |  | 758.5, 0.175, -55.67 ± 371.64 | 117.5, 0.370, 66.67 ± 407.18 |

**No mOTU <10 Specimens**

| Range measures | Month | Value | Phoridae | Mycetophilidae |
| --- | --- | --- | --- | --- |
| Highest elevation | March to April | V; p-value; mean with SD | 116.5, 0.001**, 192.59 ± 263.01 | - |
|  | April to May |  | 0, <0.001***, 379.31 ± 346.79 | - |
|  | May to June |  | 722, 0.165, 43.18 ± 342.34 | - |
|  | June to July |  | 818.5, 0.612, -10.94 ± 306.68 | 33，0.037*, 181.82 ± 227.24 |
|  | July to August |  | 1161.5, 0.035*, 46.27 ± 288.02 | 66, 0.003**, 147.83 ± 183.08 |
|  | August to September |  | 935, 0.093, -44.64 ± 305.75 | 106, 0.035*, -62.5 ± 156.06 |
|  | September to October |  | 258, 0.002**, -124.39 ± 356.44 | 12, 0.407, -50 ± 200 |
| Weighted mean elevation | March to April | V; p-value; mean with SD | 264, 0.001**, 117.59 ± 170.01 | - |
|  | April to May |  | 249, <0.001***, 147.68 ± 217.01 | - |
|  | May to June |  | 2845.5, <0.001***, 145.75 ± 249.1 | - |
|  | June to July |  | 2682, 0.050, -34.62 ± 205.92 | 51，0.019*, 146.27 ± 188.84 |
|  | July to August |  | 4713.5, 0.037*, 34.98 ± 226.26 | 92.5, 0.654, -3.52 ± 119.8 |
|  | August to September |  | 2341, 0.341, 32.65 ± 231.17 | 191.5, 0.405, 19.12 ± 123.69 |
|  | September to October |  | 596.5, <0.001***, -131.58 ± 276.06 | 52.5, 1, -9.62 ± 171.77 |
| Lowest elevation | March to April | V; p-value; mean with SD | 25, 0.803, 14.81 ± 174.76 | - |
|  | April to May |  | 225.5, 0.372, -31.03 ± 267.02 | - |
|  | May to June |  | 792.5, <0.001***, 129.55 ± 318.1 | - |
|  | June to July |  | 818.5, 0.612, -34.38 ± 252.37 | 27，0.626, 54.55 ± 284.13 |
|  | July to August |  | 1528.5, 0.44, 19.4 ± 299.24 | 2.5，<0.001***, -260.87 ± 244.4 |
|  | August to September |  | 750.5, 0.022*, 89.29 ± 375.69 | 30, 0.008**, 150 ± 291.82 |
|  | September to October |  | 549.5, 0.390, -36.59 ± 354.34 | 46, 0.606, 50 ± 441.21 |

**Supplemental Table S10.** Results of PERMANOVAs of the differences in community composition of phorids and mycetophilids.

(a) The results of overall differences of community composition:

| Taxon | Source | Df | Sums of squares | Mean squares | F.Model | Variation (R^2^) | Pr (>F) |
| --- | --- | --- | --- | --- | --- | --- | --- |
| Phoridae | elevation | 5 | 2.028 | 0.406 | 1.020 | 0.111 | 0.397 |
|  | month | 8 | 6.245 | 0.781 | 2.455 | 0.341 | <0.001 |
| Mycetophilidae | elevation | 5 | 3.053 | 0.611 | 2.107 | 0.413 | <0.001 |
|  | month | 4 | 2.223 | 0.556 | 1.718 | 0.300 | 0.002 |

(b) The results of community pairwise comparisons between months of phorids:

| Month | distance | Df | Sums of squares | F.Model | Variation (R^2^) | Pr (>F) | P_adj_BH |
| --- | --- | --- | --- | --- | --- | --- | --- |
| JUL/MAY | Bray-Curtis | 1 | 0.469 | 1.514 | 0.131 | 0.079 | 0.095 |
| JUL/OCT | Bray-Curtis | 1 | 0.963 | 3.125 | 0.258 | 0.002 | 0.010 |
| JUL/APR | Bray-Curtis | 1 | 1.192 | 4.187 | 0.295 | 0.004 | 0.010 |
| JUL/SEP | Bray-Curtis | 1 | 0.761 | 2.460 | 0.215 | 0.005 | 0.011 |
| JUL/JUN | Bray-Curtis | 1 | 0.308 | 1.023 | 0.093 | 0.465 | 0.465 |
| JUL/AUG | Bray-Curtis | 1 | 0.354 | 1.172 | 0.105 | 0.266 | 0.282 |
| JUL/MAR | Bray-Curtis | 1 | 1.351 | 4.886 | 0.352 | 0.003 | 0.010 |
| JUL/NOV | Bray-Curtis | 1 | 0.681 | 2.207 | 0.269 | 0.044 | 0.059 |
| MAY/OCT | Bray-Curtis | 1 | 0.850 | 2.555 | 0.221 | 0.003 | 0.010 |
| MAY/APR | Bray-Curtis | 1 | 0.819 | 2.668 | 0.211 | 0.005 | 0.011 |
| MAY/SEP | Bray-Curtis | 1 | 0.813 | 2.434 | 0.213 | 0.004 | 0.010 |
| MAY/JUN | Bray-Curtis | 1 | 0.395 | 1.220 | 0.109 | 0.232 | 0.253 |
| MAY/AUG | Bray-Curtis | 1 | 0.584 | 1.797 | 0.152 | 0.006 | 0.012 |
| MAY/MAR | Bray-Curtis | 1 | 1.149 | 3.815 | 0.298 | 0.003 | 0.010 |
| MAY/NOV | Bray-Curtis | 1 | 0.629 | 1.820 | 0.233 | 0.029 | 0.045 |
| OCT/APR | Bray-Curtis | 1 | 1.173 | 3.854 | 0.300 | 0.006 | 0.012 |
| OCT/SEP | Bray-Curtis | 1 | 0.395 | 1.180 | 0.128 | 0.22 | 0.248 |
| OCT/JUN | Bray-Curtis | 1 | 0.986 | 3.051 | 0.253 | 0.003 | 0.010 |
| OCT/AUG | Bray-Curtis | 1 | 0.874 | 2.695 | 0.230 | 0.004 | 0.010 |
| OCT/MAR | Bray-Curtis | 1 | 1.221 | 4.102 | 0.339 | 0.009 | 0.017 |
| OCT/NOV | Bray-Curtis | 1 | 0.454 | 1.302 | 0.207 | 0.134 | 0.156 |
| APR/SEP | Bray-Curtis | 1 | 1.147 | 3.751 | 0.294 | 0.002 | 0.010 |
| APR/JUN | Bray-Curtis | 1 | 1.166 | 3.908 | 0.281 | 0.002 | 0.010 |
| APR/AUG | Bray-Curtis | 1 | 1.039 | 3.473 | 0.258 | 0.003 | 0.010 |
| APR/MAR | Bray-Curtis | 1 | 0.264 | 0.967 | 0.097 | 0.428 | 0.440 |
| APR/NOV | Bray-Curtis | 1 | 0.744 | 2.457 | 0.291 | 0.037 | 0.053 |
| SEP/JUN | Bray-Curtis | 1 | 0.831 | 2.562 | 0.222 | 0.002 | 0.010 |
| SEP/AUG | Bray-Curtis | 1 | 0.537 | 1.651 | 0.155 | 0.033 | 0.050 |
| SEP/MAR | Bray-Curtis | 1 | 1.200 | 4.013 | 0.334 | 0.01 | 0.018 |
| SEP/NOV | Bray-Curtis | 1 | 0.547 | 1.558 | 0.238 | 0.047 | 0.060 |
| JUN/AUG | Bray-Curtis | 1 | 0.554 | 1.754 | 0.149 | 0.029 | 0.045 |
| JUN/MAR | Bray-Curtis | 1 | 1.297 | 4.449 | 0.331 | 0.004 | 0.010 |
| JUN/NOV | Bray-Curtis | 1 | 0.671 | 2.027 | 0.252 | 0.044 | 0.059 |
| AUG/MAR | Bray-Curtis | 1 | 1.225 | 4.187 | 0.318 | 0.001 | 0.010 |
| AUG/NOV | Bray-Curtis | 1 | 0.655 | 1.970 | 0.247 | 0.028 | 0.045 |
| MAR/NOV | Bray-Curtis | 1 | 0.785 | 2.690 | 0.350 | 0.056 | 0.070 |

(c) The results of community pairwise comparisons between elevations of mycetophilids:

| Elevation  (m a.s.l.) | distance | Df | Sums of squares | F.Model | Variation (R^2^) | Pr (>F) | P_adj_BH |
| --- | --- | --- | --- | --- | --- | --- | --- |
| 1200/1400 | Bray-Curtis | 1 | 0.604 | 2.147 | 0.263 | 0.038 | 0.125 |
| 1200/1600 | Bray-Curtis | 1 | 0.602 | 1.898 | 0.192 | 0.046 | 0.125 |
| 1200/1800 | Bray-Curtis | 1 | 0.652 | 1.919 | 0.215 | 0.022 | 0.125 |
| 1200/1000 | Bray-Curtis | 1 | 0.502 | 1.574 | 0.239 | 0.096 | 0.125 |
| 1200/800 | Bray-Curtis | 1 | 0.487 | 1.628 | 0.246 | 0.092 | 0.125 |
| 1400/1600 | Bray-Curtis | 1 | 0.777 | 3.004 | 0.334 | 0.018 | 0.125 |
| 1400/1800 | Bray-Curtis | 1 | 0.777 | 2.787 | 0.358 | 0.031 | 0.125 |
| 1400/1000 | Bray-Curtis | 1 | 0.792 | 3.888 | 0.564 | 0.100 | 0.125 |
| 1400/800 | Bray-Curtis | 1 | 0.523 | 3.072 | 0.506 | 0.100 | 0.125 |
| 1600/1800 | Bray-Curtis | 1 | 0.604 | 1.888 | 0.212 | 0.058 | 0.125 |
| 1600/1000 | Bray-Curtis | 1 | 0.697 | 2.391 | 0.324 | 0.097 | 0.125 |
| 1600/800 | Bray-Curtis | 1 | 0.700 | 2.579 | 0.340 | 0.089 | 0.125 |
| 1800/1000 | Bray-Curtis | 1 | 0.495 | 1.524 | 0.276 | 0.133 | 0.143 |
| 1800/800 | Bray-Curtis | 1 | 0.529 | 1.766 | 0.306 | 0.133 | 0.143 |
| 1000/800 | Bray-Curtis | 1 | 0.220 | 1.056 | 0.345 | 0.667 | 0.667 |

**Supplemental Table S11.** Beta diversity of Phoridae and Mycetophilidae across (a) elevations and (b) months using trimmed datasets and with samples < 10 specimens removed. Multiple-site dissimilarity values and their partitions (turnover and nestedness) are indicated at the top of each table. Pairwise turnover values are above and pairwise nestedness values are below the diagonal.

(a)

| **No singletons** | |  |  |  |  |  |  |
| --- | --- | --- | --- | --- | --- | --- | --- |
| Phoridae | Overall Dissimilarity:0.807 | | | Overall Turnover:0.680 | | Overall Nestedness:0.127 | |
|  | Elevation  (m a.s.l.) | 800 | 1000 | 1200 | 1400 | 1600 | 1800 |
|  | 800 | 0 | 0.202 | 0.524 | 0.322 | 0.612 | 0.745 |
|  | 1000 | 0.327 | 0 | 0.573 | 0.211 | 0.714 | 0.577 |
|  | 1200 | 0.077 | 0.114 | 0 | 0.190 | 0.602 | 0.641 |
|  | 1400 | 0.351 | 0.603 | 0.507 | 0 | 0.349 | 0.606 |
|  | 1600 | 0.094 | 0.053 | 0.034 | 0.440 | 0 | 0.423 |
|  | 1800 | 0.017 | 0.196 | 0.080 | 0.184 | 0.175 | 0 |
| Mycetophilidae | Overall Dissimilarity:0.829 | | | Overall Turnover:0.711 | | Overall Nestedness:0.118 | |
|  | Elevation  (m a.s.l.) | 800 | 1000 | 1200 | 1400 | 1600 | 1800 |
|  | 800 | 0 | 0.481 | 0.246 | 0.485 | 0.413 | 0.653 |
|  | 1000 | 0.059 | 0 | 0.398 | 0.789 | 0.624 | 0.624 |
|  | 1200 | 0.411 | 0.373 | 0 | 0.642 | 0.582 | 0.500 |
|  | 1400 | 0.279 | 0.130 | 0.002 | 0 | 0.701 | 0.681 |
|  | 1600 | 0.338 | 0.243 | 0.019 | 0.015 | 0 | 0.418 |
|  | 1800 | 0.015 | 0.058 | 0.257 | 0.163 | 0.318 | 0 |

**No doubletons**

| Phoridae | Overall Dissimilarity:0.806 | | | Overall Turnover:0.678 | | Overall Nestedness:0.128 | |
| --- | --- | --- | --- | --- | --- | --- | --- |
|  | Elevation  (m a.s.l.) | 800 | 1000 | 1200 | 1400 | 1600 | 1800 |
|  | 800 | 0 | 0.196 | 0.520 | 0.320 | 0.609 | 0.742 |
|  | 1000 | 0.331 | 0 | 0.569 | 0.207 | 0.712 | 0.572 |
|  | 1200 | 0.077 | 0.116 | 0 | 0.185 | 0.600 | 0.638 |
|  | 1400 | 0.351 | 0.607 | 0.510 | 0 | 0.347 | 0.604 |
|  | 1600 | 0.095 | 0.054 | 0.034 | 0.441 | 0 | 0.418 |
|  | 1800 | 0.017 | 0.199 | 0.081 | 0.185 | 0.177 | 0 |
| Mycetophilidae | Overall Dissimilarity:0.828 | | | Overall Turnover:0.706 | | Overall Nestedness:0.122 | |
|  | Elevation  (m a.s.l.) | 800 | 1000 | 1200 | 1400 | 1600 | 1800 |
|  | 800 | 0 | 0.457 | 0.228 | 0.488 | 0.401 | 0.642 |
|  | 1000 | 0.066 | 0 | 0.370 | 0.787 | 0.614 | 0.606 |
|  | 1200 | 0.426 | 0.397 | 0 | 0.636 | 0.581 | 0.486 |
|  | 1400 | 0.279 | 0.133 | 0.004 | 0 | 0.698 | 0.672 |
|  | 1600 | 0.348 | 0.253 | 0.018 | 0.016 | 0 | 0.407 |
|  | 1800 | 0.016 | 0.065 | 0.268 | 0.168 | 0.327 | 0 |

**No mOTU <5 specimens**

| Phoridae | Overall Dissimilarity:0.806 | | Overall Turnover:0.675 | | | Overall Nestedness:0.131 | |
| --- | --- | --- | --- | --- | --- | --- | --- |
|  | Elevation  (m a.s.l.) | 800 | 1000 | 1200 | 1400 | 1600 | 1800 |
|  | 800 | 0 | 0.182 | 0.516 | 0.312 | 0.604 | 0.745 |
|  | 1000 | 0.343 | 0 | 0.565 | 0.201 | 0.709 | 0.570 |
|  | 1200 | 0.080 | 0.119 | 0 | 0.182 | 0.597 | 0.636 |
|  | 1400 | 0.355 | 0.615 | 0.513 | 0 | 0.352 | 0.610 |
|  | 1600 | 0.098 | 0.056 | 0.035 | 0.439 | 0 | 0.416 |
|  | 1800 | 0.016 | 0.202 | 0.082 | 0.183 | 0.178 | 0 |
| Mycetophilidae | Overall Dissimilarity:0.832 | | Overall Turnover:0.704 | | | Overall Nestedness:0.128 | |
|  | Elevation  (m a.s.l.) | 800 | 1000 | 1200 | 1400 | 1600 | 1800 |
|  | 800 | 0 | 0.406 | 0.197 | 0.486 | 0.387 | 0.620 |
|  | 1000 | 0.086 | 0 | 0.368 | 0.830 | 0.632 | 0.585 |
|  | 1200 | 0.472 | 0.427 | 0 | 0.638 | 0.580 | 0.488 |
|  | 1400 | 0.297 | 0.113 | 0.005 | 0 | 0.706 | 0.690 |
|  | 1600 | 0.375 | 0.256 | 0.016 | 0.015 | 0 | 0.429 |
|  | 1800 | 0.032 | 0.094 | 0.271 | 0.161 | 0.318 | 0 |

**No mOTU <10 specimens**

| Phoridae | Overall Dissimilarity:0.804 | | | Overall Turnover:0.665 | | Overall Nestedness:0.138 | |
| --- | --- | --- | --- | --- | --- | --- | --- |
|  | Elevation (m a.s.l.) | 800 | 1000 | 1200 | 1400 | 1600 | 1800 |
|  | 800 | 0 | 0.162 | 0.514 | 0.295 | 0.594 | 0.738 |
|  | 1000 | 0.363 | 0 | 0.549 | 0.177 | 0.703 | 0.551 |
|  | 1200 | 0.076 | 0.133 | 0 | 0.171 | 0.586 | 0.621 |
|  | 1400 | 0.365 | 0.639 | 0.517 | 0 | 0.344 | 0.603 |
|  | 1600 | 0.103 | 0.060 | 0.041 | 0.446 | 0 | 0.392 |
|  | 1800 | 0.018 | 0.218 | 0.084 | 0.185 | 0.192 | 0 |
| Mycetophilidae | Overall Dissimilarity:0.832 | | | Overall Turnover:0.702 | | Overall Nestedness:0.130 | |
|  | Elevation (m a.s.l.) | 800 | 1000 | 1200 | 1400 | 1600 | 1800 |
|  | 800 | 0 | 0.370 | 0.160 | 0.481 | 0.405 | 0.603 |
|  | 1000 | 0.085 | 0 | 0.340 | 0.820 | 0.630 | 0.570 |
|  | 1200 | 0.501 | 0.447 | 0 | 0.633 | 0.578 | 0.486 |
|  | 1400 | 0.306 | 0.121 | 0.004 | 0 | 0.710 | 0.676 |
|  | 1600 | 0.365 | 0.255 | 0.010 | 0.010 | 0 | 0.419 |
|  | 1800 | 0.024 | 0.083 | 0.286 | 0.178 | 0.333 | 0 |

(b)

**No singletons**

| Phoridae | Overall Dissimilarity:0.870 | | | | Overall Turnover:0.780 | | | Overall Nestedness:0.090 | | |
| --- | --- | --- | --- | --- | --- | --- | --- | --- | --- | --- |
|  | Month | MAR | APR | MAY | JUN | JUL | AUG | SEP | OCT | NOV |
|  | MAR | 0 | 0.332 | 0.902 | 0.927 | 0.927 | 0.877 | 0.927 | 0.936 | 0.431 |
|  | APR | 0.178 | 0 | 0.587 | 0.714 | 0.756 | 0.671 | 0.900 | 0.910 | 0.611 |
|  | MAY | 0.000 | 0.111 | 0 | 0.297 | 0.446 | 0.558 | 0.779 | 0.800 | 0.417 |
|  | JUN | 0.024 | 0.157 | 0.231 | 0 | 0.260 | 0.587 | 0.695 | 0.724 | 0.194 |
|  | JUL | 0.007 | 0.086 | 0.051 | 0.181 | 0 | 0.485 | 0.677 | 0.762 | 0.236 |
|  | AUG | 0.025 | 0.147 | 0.090 | 0.056 | 0.059 | 0 | 0.401 | 0.654 | 0.347 |
|  | SEP | 0.015 | 0.007 | 0.046 | 0.153 | 0.094 | 0.235 | 0 | 0.516 | 0.292 |
|  | OCT | 0.022 | 0.007 | 0.068 | 0.167 | 0.100 | 0.176 | 0.071 | 0 | 0.250 |
|  | NOV | 0.530 | 0.344 | 0.543 | 0.777 | 0.720 | 0.623 | 0.636 | 0.648 | 0 |
| Mycetophilidae | Overall Dissimilarity:0.792 | | | | Overall Turnover:0.371 | | | Overall Nestedness:0.421 | | |
|  | Month |  | | | JUN | JUL | AUG | SEP | OCT |  |
|  | JUN |  | | | 0 | 0.507 | 0.187 | 0.360 | 0.533 |  |
|  | JUL |  |  |  | 0.194 | 0 | 0.110 | 0.221 | 0.822 |  |
|  | AUG |  |  |  | 0.721 | 0.674 | 0 | 0.092 | 0.420 |  |
|  | SEP |  |  |  | 0.490 | 0.415 | 0.342 | 0 | 0.385 |  |
|  | OCT |  |  |  | 0.180 | 0.002 | 0.442 | 0.332 | 0 |  |

**No doubletons**

| Phoridae | Overall Dissimilarity:0.870 | | | | Overall Turnover:0.779 | | | Overall Nestedness:0.090 | | |
| --- | --- | --- | --- | --- | --- | --- | --- | --- | --- | --- |
|  | Month | MAR | APR | MAY | JUN | JUL | AUG | SEP | OCT | NOV |
|  | MAR | 0 | 0.332 | 0.902 | 0.927 | 0.927 | 0.877 | 0.926 | 0.936 | 0.414 |
|  | APR | 0.179 | 0 | 0.587 | 0.713 | 0.755 | 0.670 | 0.899 | 0.909 | 0.600 |
|  | MAY | 0.000 | 0.110 | 0 | 0.294 | 0.445 | 0.557 | 0.777 | 0.798 | 0.400 |
|  | JUN | 0.024 | 0.157 | 0.231 | 0 | 0.255 | 0.585 | 0.694 | 0.721 | 0.171 |
|  | JUL | 0.006 | 0.085 | 0.049 | 0.184 | 0 | 0.481 | 0.675 | 0.760 | 0.229 |
|  | AUG | 0.025 | 0.147 | 0.089 | 0.056 | 0.059 | 0 | 0.396 | 0.650 | 0.329 |
|  | SEP | 0.016 | 0.006 | 0.047 | 0.154 | 0.096 | 0.239 | 0 | 0.512 | 0.271 |
|  | OCT | 0.022 | 0.008 | 0.070 | 0.168 | 0.101 | 0.179 | 0.071 | 0 | 0.243 |
|  | NOV | 0.546 | 0.355 | 0.560 | 0.800 | 0.728 | 0.641 | 0.655 | 0.656 | 0 |
| Mycetophilidae | Overall Dissimilarity:0.790 | | | | Overall Turnover:0.362 | | | Overall Nestedness:0.428 | | |
|  | Month |  | | | JUN | JUL | AUG | SEP | OCT |  |
|  | JUN |  | | | 0 | 0.500 | 0.181 | 0.333 | 0.514 |  |
|  | JUL |  |  |  | 0.196 | 0 | 0.091 | 0.194 | 0.818 |  |
|  | AUG |  |  |  | 0.728 | 0.692 | 0 | 0.091 | 0.423 |  |
|  | SEP |  |  |  | 0.515 | 0.439 | 0.337 | 0 | 0.381 |  |
|  | OCT |  |  |  | 0.194 | 0.002 | 0.438 | 0.333 | 0 |  |

**No mOTU <5 specimens**

| Phoridae | Overall Dissimilarity:0.870 | | | | Overall Turnover:0.778 | | | Overall Nestedness:0.092 | | |
| --- | --- | --- | --- | --- | --- | --- | --- | --- | --- | --- |
|  | Month | MAR | APR | MAY | JUN | JUL | AUG | SEP | OCT | NOV |
|  | MAR | 0 | 0.331 | 0.900 | 0.927 | 0.927 | 0.877 | 0.924 | 0.934 | 0.465 |
|  | APR | 0.179 | 0 | 0.586 | 0.712 | 0.754 | 0.669 | 0.900 | 0.908 | 0.651 |
|  | MAY | 0.000 | 0.109 | 0 | 0.287 | 0.438 | 0.553 | 0.775 | 0.796 | 0.395 |
|  | JUN | 0.024 | 0.157 | 0.234 | 0 | 0.249 | 0.580 | 0.689 | 0.718 | 0.163 |
|  | JUL | 0.006 | 0.084 | 0.050 | 0.186 | 0 | 0.481 | 0.668 | 0.758 | 0.116 |
|  | AUG | 0.024 | 0.145 | 0.089 | 0.058 | 0.058 | 0 | 0.389 | 0.644 | 0.209 |
|  | SEP | 0.017 | 0.005 | 0.049 | 0.158 | 0.100 | 0.244 | 0 | 0.509 | 0.279 |
|  | OCT | 0.023 | 0.009 | 0.071 | 0.171 | 0.103 | 0.182 | 0.070 | 0 | 0.256 |
|  | NOV | 0.512 | 0.324 | 0.579 | 0.819 | 0.852 | 0.768 | 0.674 | 0.680 | 0 |
| Mycetophilidae | Overall Dissimilarity:0.795 | | | | Overall Turnover:0.344 | | | Overall Nestedness:0.451 | | |
|  | Month |  | | | JUN | JUL | AUG | SEP | OCT |  |
|  | JUN |  | | | 0 | 0.431 | 0.190 | 0.259 | 0.500 |  |
|  | JUL |  |  |  | 0.250 | 0 | 0.094 | 0.161 | 0.864 |  |
|  | AUG |  |  |  | 0.734 | 0.703 | 0 | 0.084 | 0.456 |  |
|  | SEP |  |  |  | 0.599 | 0.480 | 0.335 | 0 | 0.381 |  |
|  | OCT |  |  |  | 0.217 | 0.001 | 0.424 | 0.357 | 0 |  |

**No mOTU <10 specimens**

| Phoridae | Overall Dissimilarity:0.868 | | | | Overall Turnover:0.776 | | | Overall Nestedness:0.093 | | |
| --- | --- | --- | --- | --- | --- | --- | --- | --- | --- | --- |
|  | Month | MAR | APR | MAY | JUN | JUL | AUG | SEP | OCT | NOV |
|  | MAR | 0 | 0.326 | 0.897 | 0.928 | 0.928 | 0.878 | 0.919 | 0.933 | 0.463 |
|  | APR | 0.182 | 0 | 0.588 | 0.714 | 0.756 | 0.670 | 0.902 | 0.909 | 0.659 |
|  | MAY | 0.002 | 0.103 | 0 | 0.269 | 0.425 | 0.543 | 0.767 | 0.793 | 0.366 |
|  | JUN | 0.023 | 0.155 | 0.246 | 0 | 0.218 | 0.565 | 0.678 | 0.710 | 0.171 |
|  | JUL | 0.004 | 0.078 | 0.045 | 0.208 | 0 | 0.478 | 0.664 | 0.755 | 0.098 |
|  | AUG | 0.021 | 0.139 | 0.088 | 0.067 | 0.060 | 0 | 0.374 | 0.640 | 0.195 |
|  | SEP | 0.020 | 0.002 | 0.053 | 0.169 | 0.101 | 0.251 | 0 | 0.511 | 0.268 |
|  | OCT | 0.025 | 0.010 | 0.072 | 0.178 | 0.102 | 0.182 | 0.065 | 0 | 0.268 |
|  | NOV | 0.515 | 0.318 | 0.608 | 0.812 | 0.870 | 0.782 | 0.684 | 0669 | 0 |
| Mycetophilidae | Overall Dissimilarity:0.792 | | | | Overall Turnover:0.324 | | | Overall Nestedness:0.468 | | |
|  | Month |  | | | JUN | JUL | AUG | SEP | OCT |  |
|  | JUN |  | | | 0 | 0.446 | 0.179 | 0.232 | 0.554 |  |
|  | JUL |  |  |  | 0.237 | 0 | 0.064 | 0.107 | 0.908 |  |
|  | AUG |  |  |  | 0.743 | 0.727 | 0 | 0.041 | 0.485 |  |
|  | SEP |  |  |  | 0.615 | 0.507 | 0.359 | 0 | 0.423 |  |
|  | OCT |  |  |  | 0.178 | 0.003 | 0.408 | 0.342 | 0 |  |

**Supplemental** **Table S12.** The results for NTI in both taxa by (a) elevation and (b) month.

(a)

|  | Elevation  (m a.s.l.) | ntaxa | mntd.obs | mntd.rand.mean | mntd.rand.sd | mntd.obs.rank | mntd.obs.p | runs | NTI |
| --- | --- | --- | --- | --- | --- | --- | --- | --- | --- |
| Phoridae | 800 | 223 | 0.878 | 0.978 | 0.049 | 15 | 0.015 | 999 | 2.033 |
|  | 1000 | 231 | 1.012 | 0.970 | 0.048 | 802 | 0.802 | 999 | -0.872 |
|  | 1200 | 276 | 0.885 | 0.928 | 0.040 | 155 | 0.155 | 999 | 1.073 |
|  | 1400 | 132 | 0.865 | 1.103 | 0.080 | 1 | 0.001 | 999 | 2.985 |
|  | 1600 | 254 | 0.747 | 0.949 | 0.043 | 1 | 0.001 | 999 | 4.701 |
|  | 1800 | 199 | 0.772 | 1.007 | 0.055 | 1 | 0.001 | 999 | 4.307 |
| Mycophilidae | 800 | 47 | 0.929 | 0.841 | 0.055 | 947 | 0.947 | 999 | -1.608 |
|  | 1000 | 31 | 0.898 | 0.917 | 0.072 | 390 | 0.39 | 999 | 0.256 |
|  | 1200 | 66 | 0.822 | 0.779 | 0.045 | 826 | 0.826 | 999 | -0.964 |
|  | 1400 | 61 | 0.803 | 0.796 | 0.047 | 559 | 0.559 | 999 | -0.161 |
|  | 1600 | 83 | 0.720 | 0.739 | 0.036 | 314 | 0.314 | 999 | 0.514 |
|  | 1800 | 42 | 0.737 | 0.864 | 0.060 | 19 | 0.019 | 999 | 2.115 |

(b)

|  | Month | ntaxa | mntd.obs | mntd.rand.mean | mntd.rand.sd | mntd.obs.rank | mntd.obs.p | runs | NTI |
| --- | --- | --- | --- | --- | --- | --- | --- | --- | --- |
| Phoridae | MAR | 62 | 0.879 | 1.274 | 0.137 | 1 | 0.001 | 999 | 2.879 |
|  | APR | 76 | 0.946 | 1.236 | 0.118 | 2 | 0.002 | 999 | 2.465 |
|  | MAY | 149 | 0.907 | 1.072 | 0.071 | 3 | 0.003 | 999 | 2.337 |
|  | JUN | 243 | 0.934 | 0.957 | 0.045 | 317 | 0.317 | 999 | 0.497 |
|  | JUL | 264 | 0.887 | 0.940 | 0.041 | 106 | 0.106 | 999 | 1.291 |
|  | AUG | 285 | 0.838 | 0.920 | 0.039 | 19 | 0.019 | 999 | 2.095 |
|  | SEP | 202 | 0.837 | 1.001 | 0.054 | 1 | 0.001 | 999 | 3.021 |
|  | OCT | 135 | 0.892 | 1.095 | 0.075 | 2 | 0.002 | 999 | 2.714 |
|  | NOV | 40 | 0.976 | 1.376 | 0.172 | 1 | 0.001 | 999 | 2.316 |
| Mycophilidae | MAY | 2 | 1.622 | 1.628 | 0.357 | 524 | 0.524 | 999 | 0.016 |
|  | JUN | 26 | 0.736 | 0.953 | 0.081 | 3 | 0.003 | 999 | 2.700 |
|  | JUL | 42 | 0.895 | 0.860 | 0.063 | 691 | 0.691 | 999 | -0.553 |
|  | AUG | 119 | 0.652 | 0.666 | 0.022 | 287 | 0.287 | 999 | 0.641 |
|  | SEP | 69 | 0.720 | 0.773 | 0.043 | 120 | 0.12 | 999 | 1.222 |
|  | OCT | 32 | 0.801 | 0.912 | 0.073 | 69 | 0.069 | 999 | 1.522 |
