## Supplementary Figure S1-S13 for "Megabarcoding reveals a tale of two very different dark taxa along the same elevational gradient"

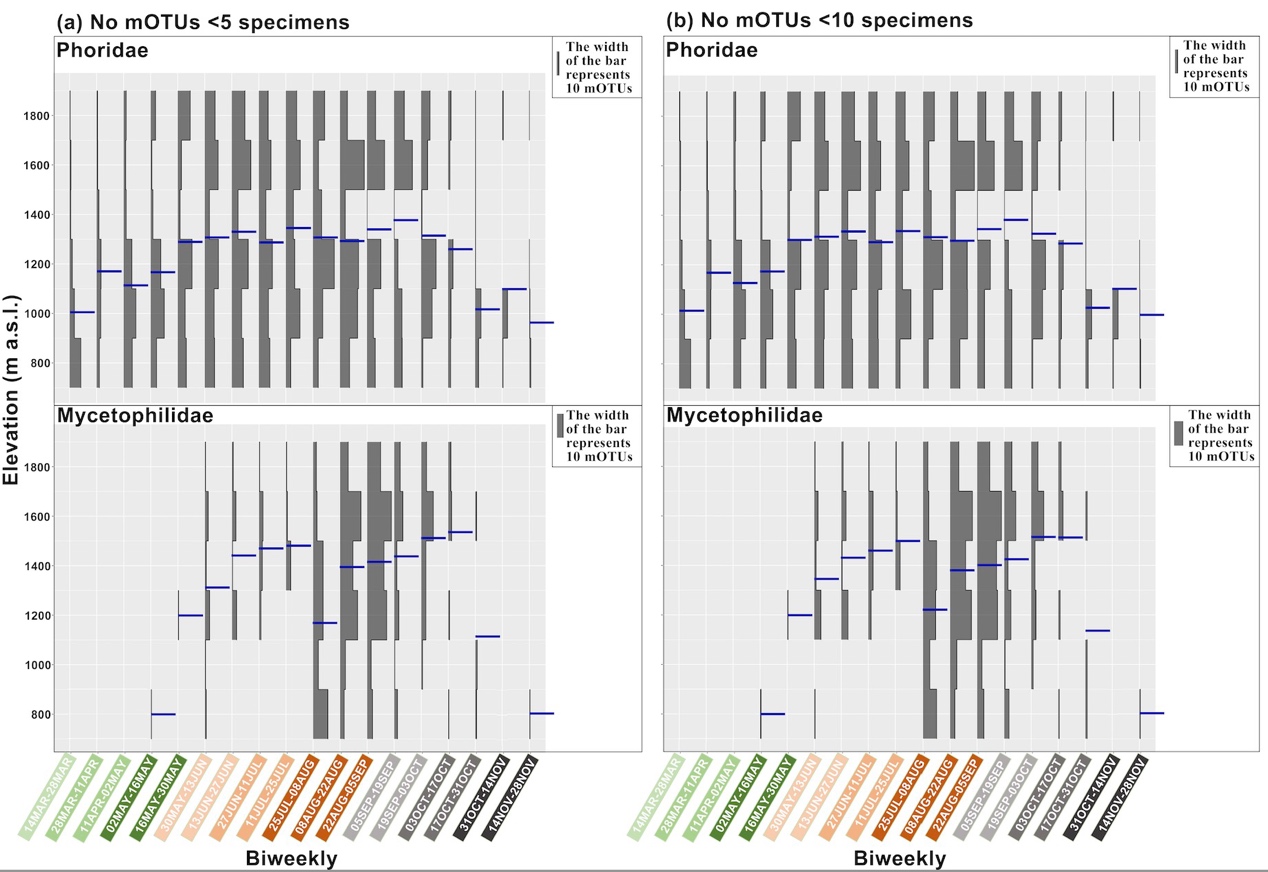

**Supplemental Figure S1.** Patterns of richness along the elevation gradient, shown in 2-week intervals, using the dataset for mOTUs with (a) less than 5 specimens and (b) less than 10 specimens removed. Blue bar indicates mean elevation per taxa at 2-week intervals (see Methods for details).

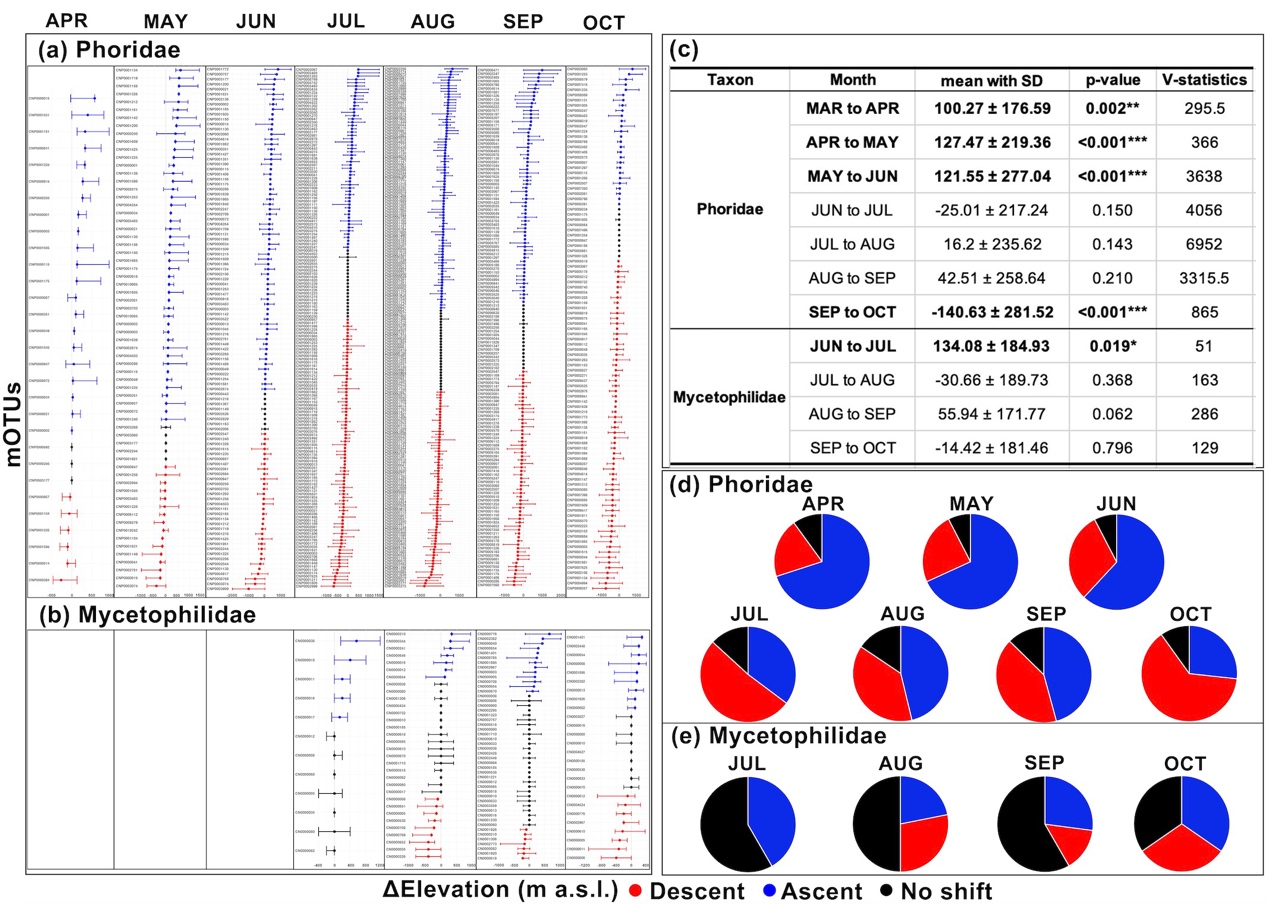

**Supplemental Figure S2.** Elevation range shifts for common mOTUs with at least one observed inter‐monthly shift (a) Phoridae and (b) Mycetophilidae. Months in the figure are in comparison to the previous month. Blue colour indicates an ascent, red colour a descent, and black indicates no change in the weighted mean elevation. Dots = weighted mean elevations; whiskers = absolute values of shift of the lowest (left) or highest (right) elevations. (c) Wilcoxon signed-rank tests for significant differences in the weighted mean elevation of consecutive months for Phoridae and Mycetophilidae (p-value: * p < 0.05; ** p < 0.01; *** p < 0.001). Proportion of ascending, descending or unchanged MOTUs for (d) Phoridae and (e) Mycetophilidae. Doubletons are removed from this dataset.

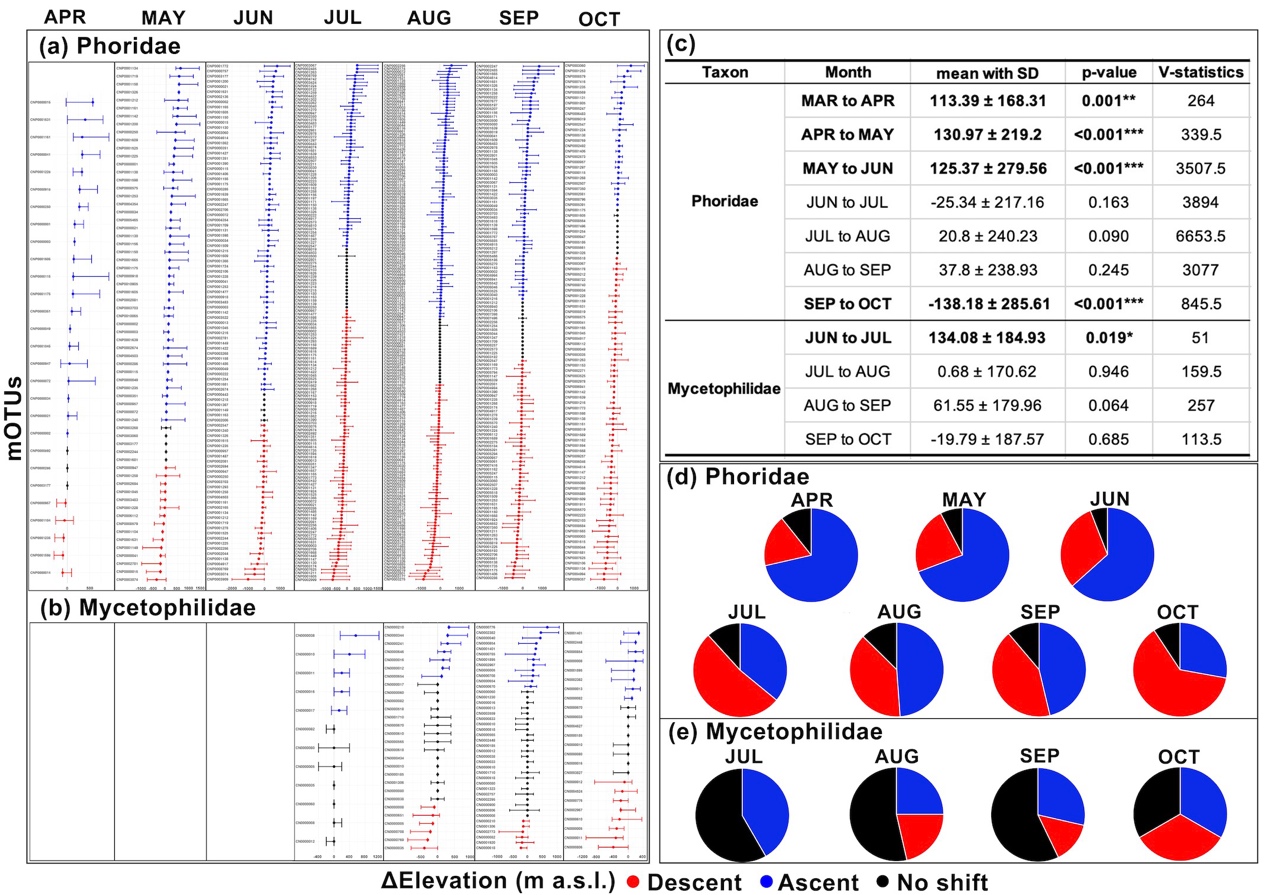

**Supplementary Figure S3.** Elevation range shifts for common mOTUs with at least one observed inter‐monthly shift (a) Phoridae and (b) Mycetophilidae. Months in the figure are in comparison to the previous month. Blue colour indicates an ascent, red colour a descent, and black indicates no change in the weighted mean elevation. Dots = weighted mean elevations; whiskers = absolute values of shift of the lowest (left) or highest (right) elevations. (c) Wilcoxon signed-rank tests for significant differences in the weighted mean elevation of consecutive months for Phoridae and Mycetophilidae (p-value: * p < 0.05; ** p < 0.01; *** p < 0.001). Proportion of ascending, descending or unchanged MOTUs for (d) Phoridae and (e) Mycetophilidae. mOTUs with less than 5 specimens are removed from this dataset.

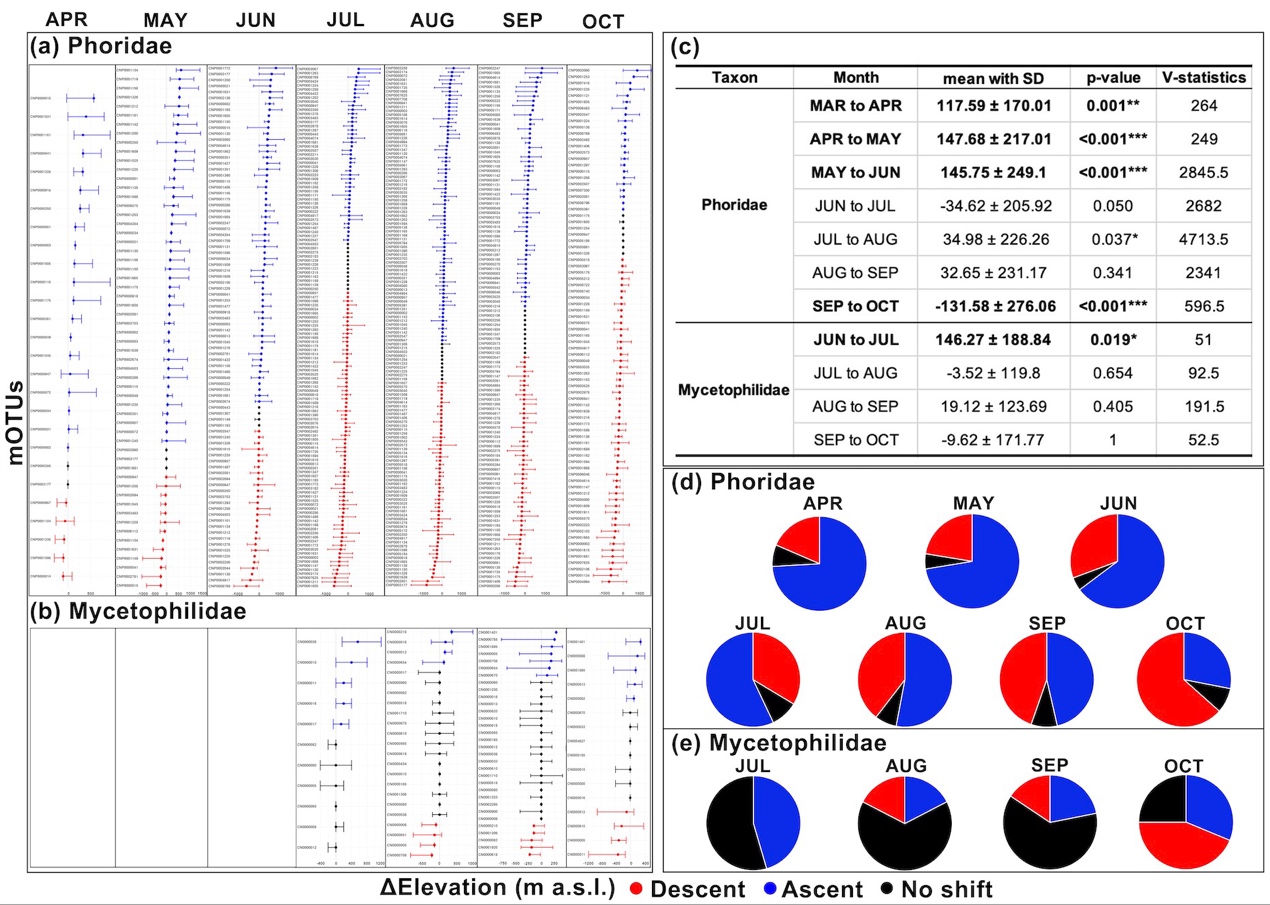

**Supplementary Figure S4.** Elevation range shifts for common mOTUs with at least one observed inter‐monthly shift (a) Phoridae and (b) Mycetophilidae. Months in the figure are in comparison to the previous month. Blue colour indicates an ascent, red colour a descent, and black indicates no change in the weighted mean elevation. Dots = weighted mean elevations; whiskers = absolute values of shift of the lowest (left) or highest (right) elevations. (c) Wilcoxon signed-rank tests for significant differences in the weighted mean elevation of consecutive months for Phoridae and Mycetophilidae (p-value: * p < 0.05; ** p < 0.01; *** p < 0.001). Proportion of ascending, descending or unchanged MOTUs for (d) Phoridae and (e) Mycetophilidae. mOTUs with less than 10 specimens are removed from this dataset.

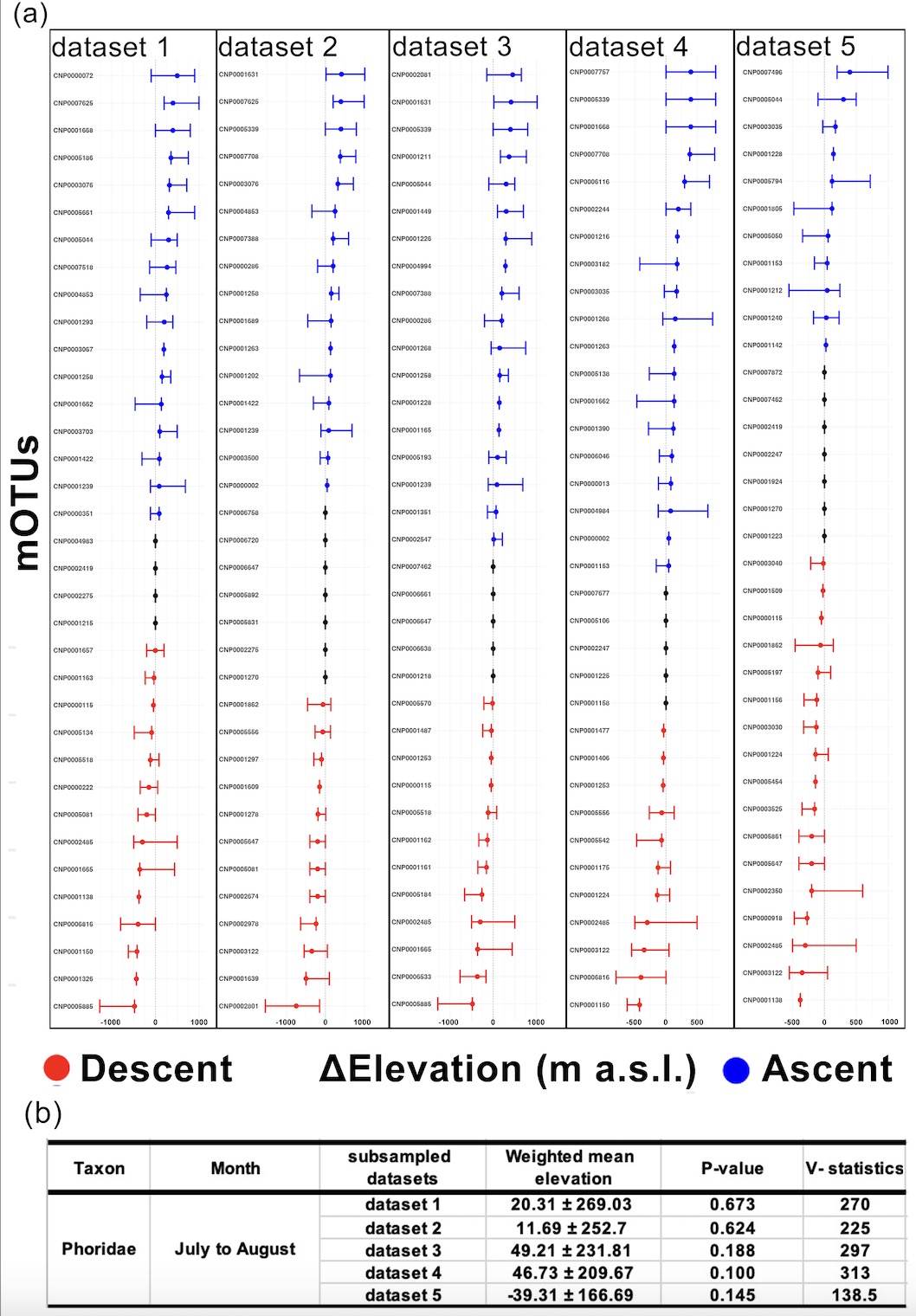

**Supplemental** **Figure S5.** From the full dataset, (a) five randomly subsampled datasets of 35 Phoridae mOTUs (ranging from 1,776 to 3,061 specimens) matching the 35 Mycetophilidae mOTUs (932 specimens) which exhibited changes in elevation from July to August. (b) Detailed results of the Wilcoxon signed-rank tests of the change in weighted mean elevation for the five subsampled datasets.

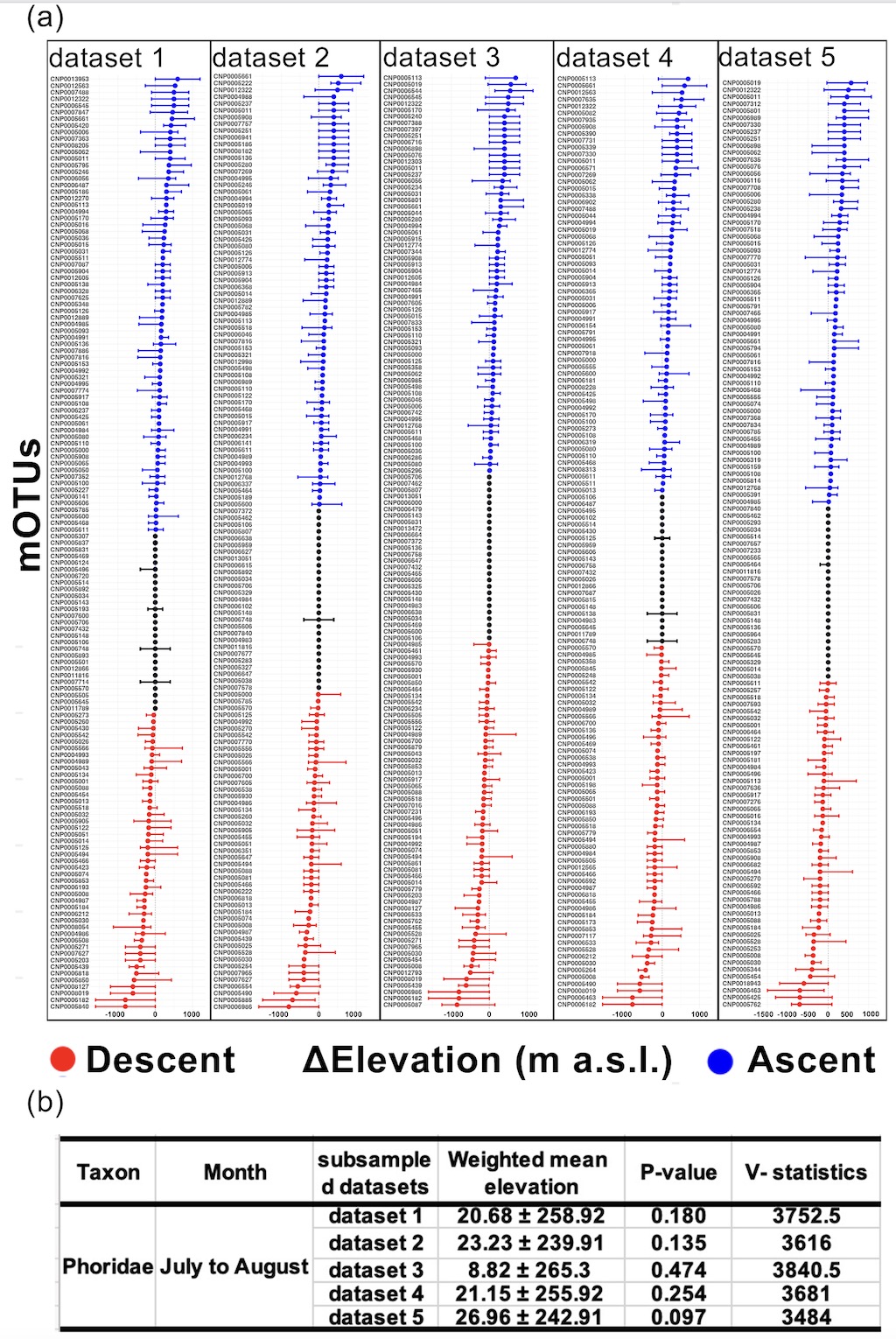

**Supplemental Figure S6.** From the full dataset, (a) five randomly subsampled datasets of 932 Phoridae specimens (ranging from 133 to 145 mOTUs) matching the 932 Mycetophilidae specimens that represented the 35 Mycetophilidae mOTUs. (b) Detailed results of the Wilcoxon signed-rank tests of the change in weighted mean elevation for the five subsampled datasets.

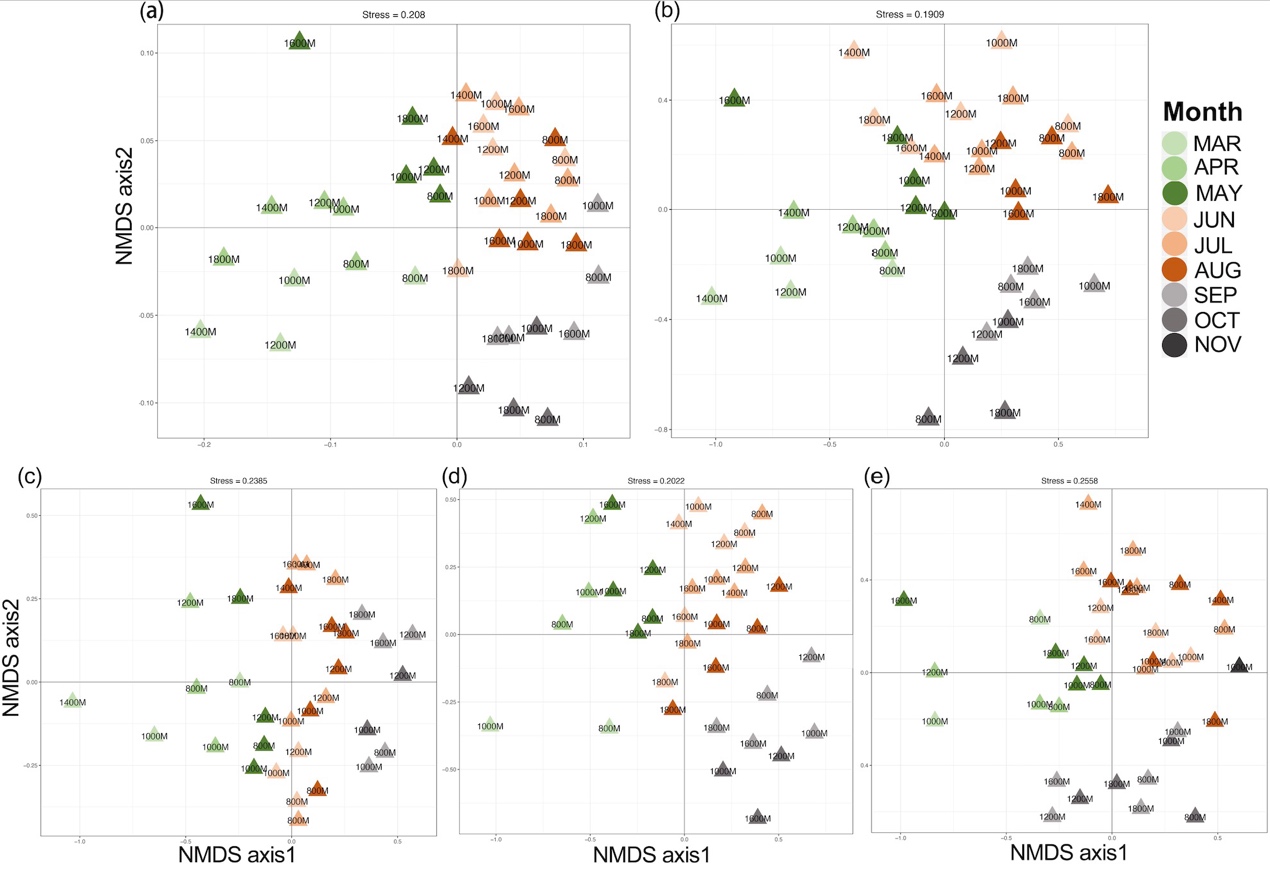

**Supplemental Figure S7.** (a)-(e) From the full dataset, five randomly subsampled datasets of 148 Phoridae mOTUs (ranging from 3,488 to 5,740 specimens) matching the 148 Mycetophilidae mOTUs were created. Non-metric Multi-dimensional Scaling (NMDS) based on Bray-Curtis dissimilarity matrices are illustrated on two-dimensional NMDS plots. Samples with less than 10 specimens are removed from these datasets.

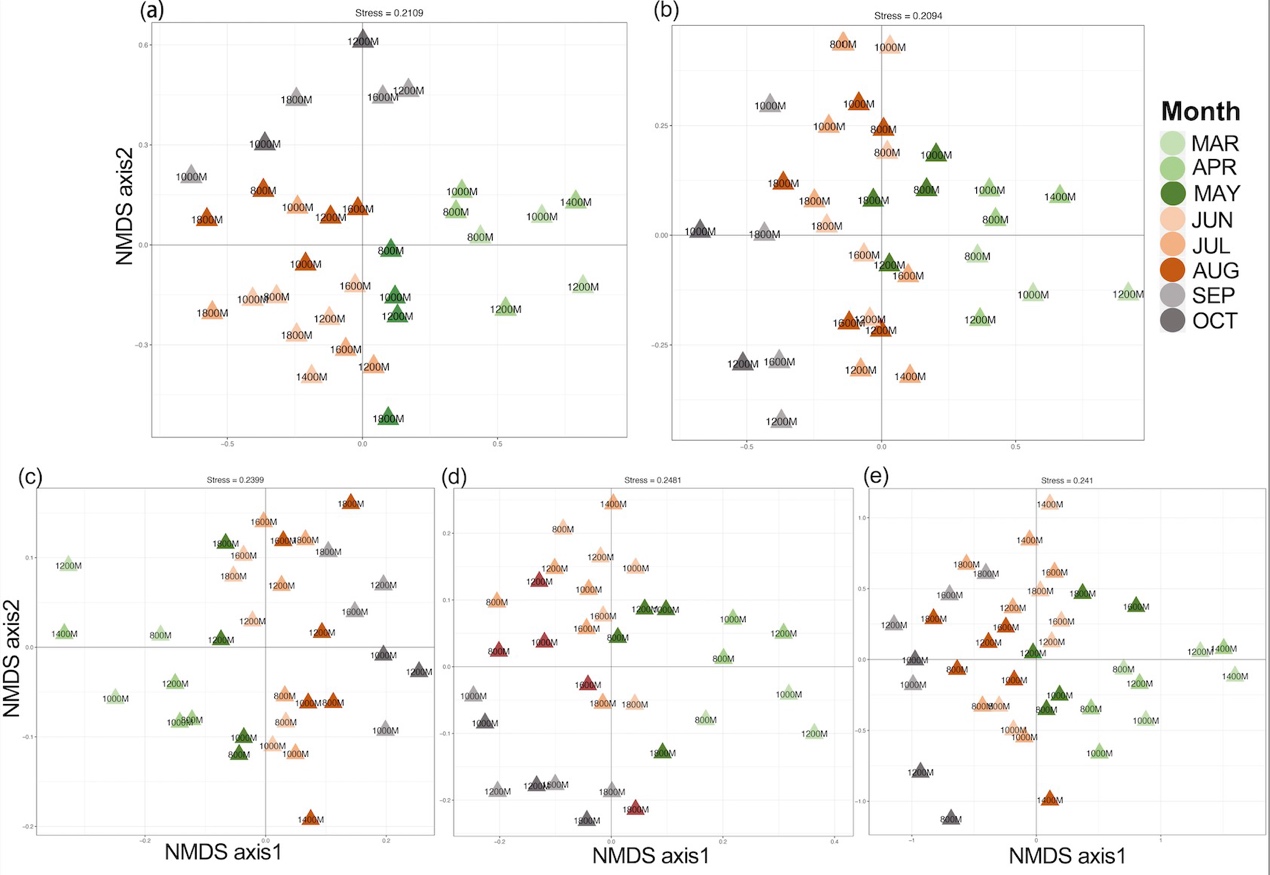

**Supplemental Figure S8.** (a)-(e) From the full dataset, five randomly subsampled datasets of 2,283 Phoridae specimens (ranging from 275 to 285 mOTUs) matching the 2,283 Mycetophilidae specimens were created. Non-metric Multi-dimensional Scaling (NMDS) based on Bray-Curtis dissimilarity matrices are illustrated on two-dimensional NMDS plots. Samples with less than 10 specimens are removed from these datasets.

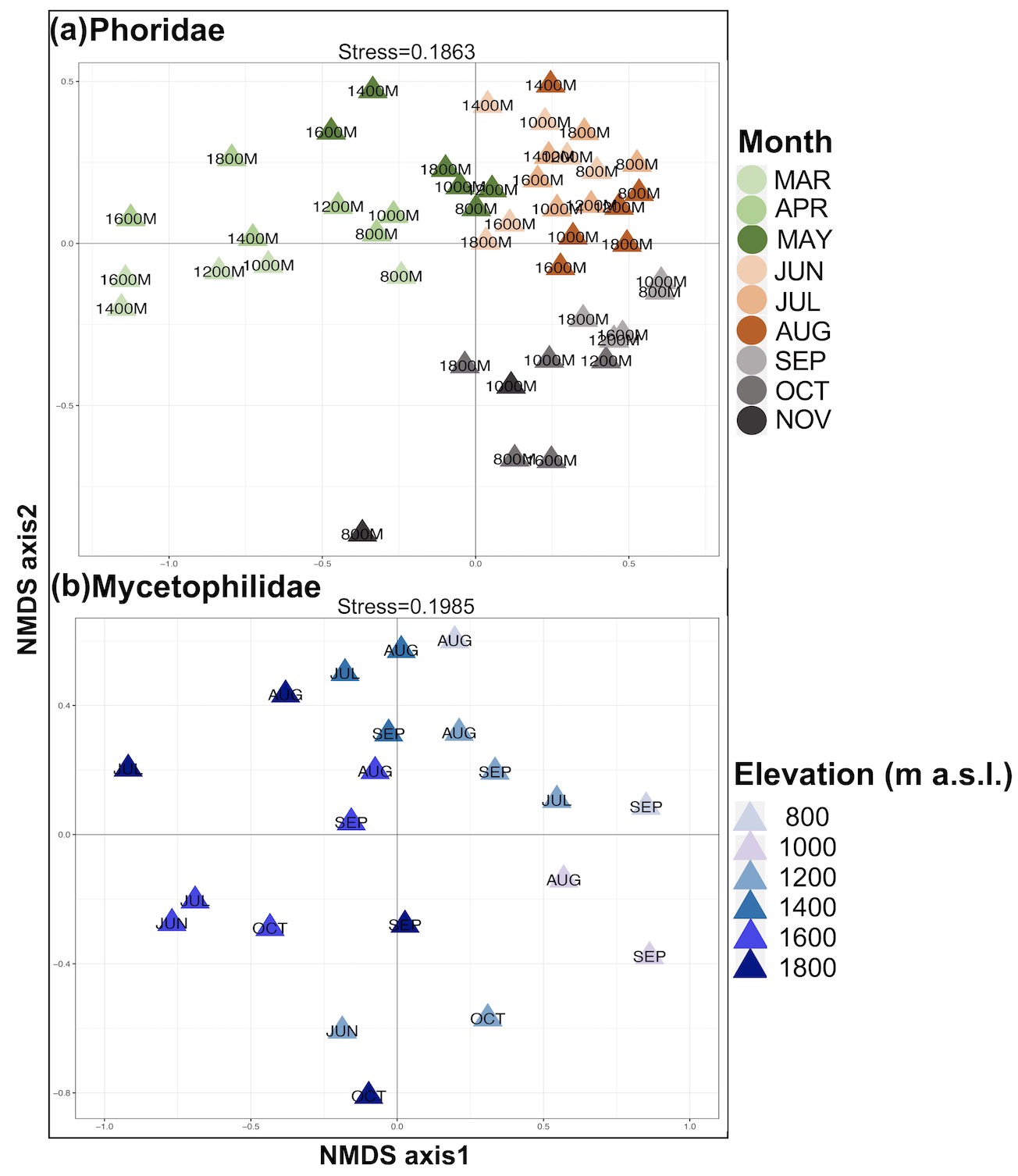

**Supplementary Figure S9**. Non-metric Multi-dimensional Scaling (NMDS) based on Bray-Curtis dissimilarity matrices are illustrated on two-dimensional NMDS plots. Singletons and samples with less than 10 specimens are removed from this dataset.

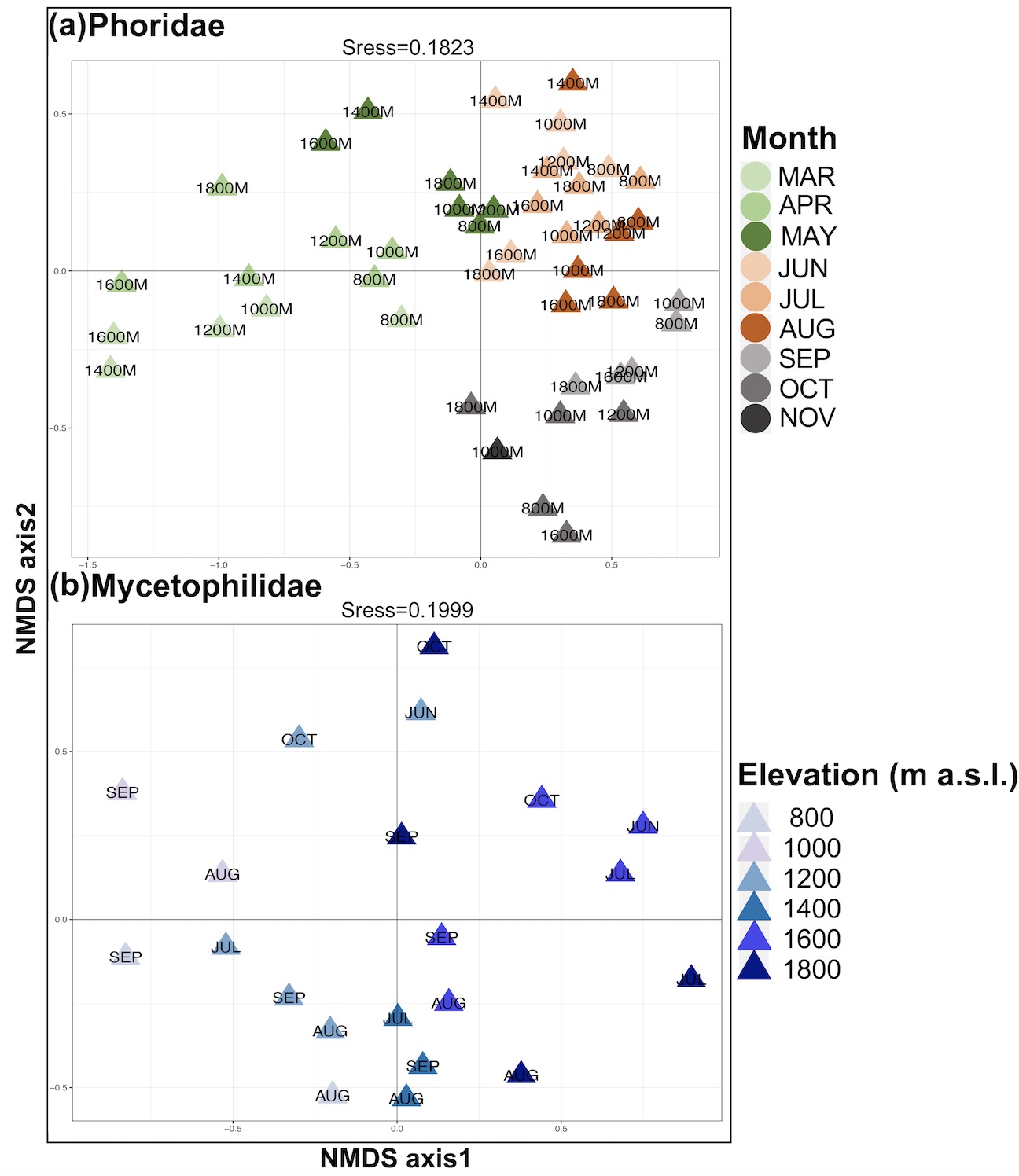

**Supplementary Figure S10.** Non-metric Multi-dimensional Scaling (NMDS) based on Bray-Curtis dissimilarity matrices are illustrated on two-dimensional NMDS plots. Doubletons and samples with less than 10 specimens are removed from this dataset.

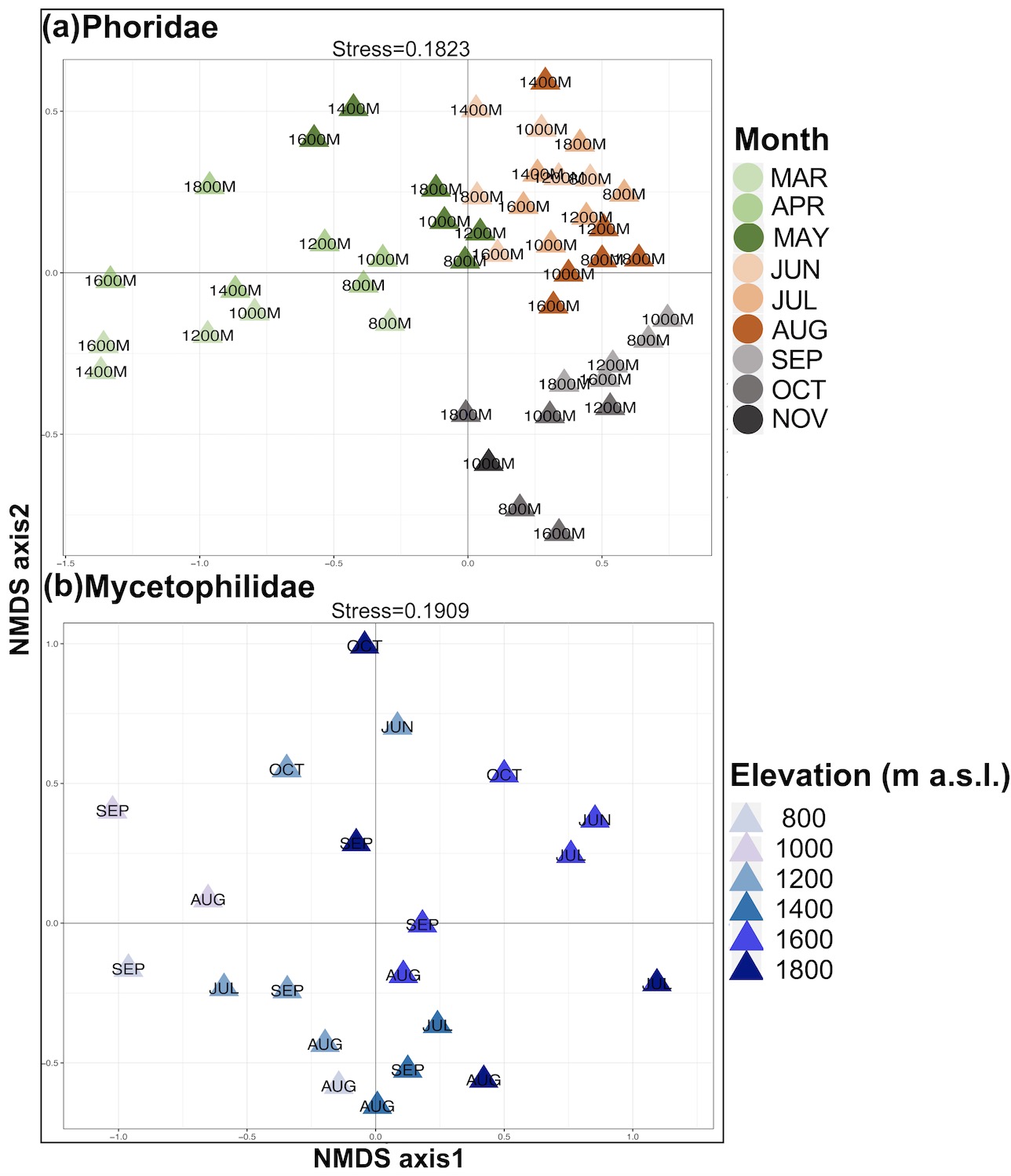

**Supplementary Figure S11.** Non-metric Multi-dimensional Scaling (NMDS) based on Bray-Curtis dissimilarity matrices are illustrated on two-dimensional NMDS plots. mOTUs with less than 5 specimens and samples with less than 10 specimens are removed from this dataset.

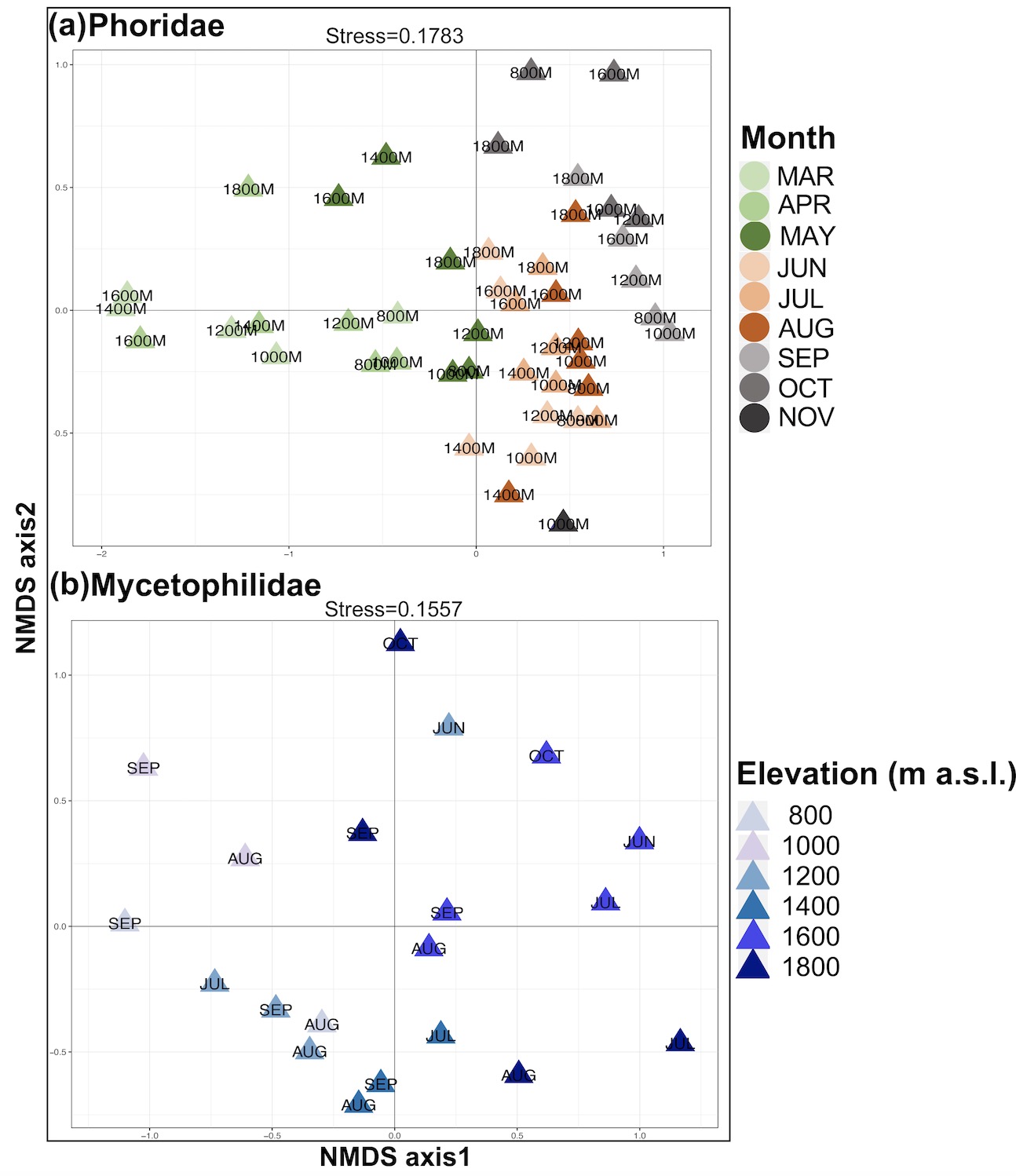

**Supplementary Figure S12.** Non-metric Multi-dimensional Scaling (NMDS) based on Bray-Curtis dissimilarity matrices are illustrated on two-dimensional NMDS plots. mOTUs with less than 10 specimens and samples with less than 10 specimens are removed from this dataset.

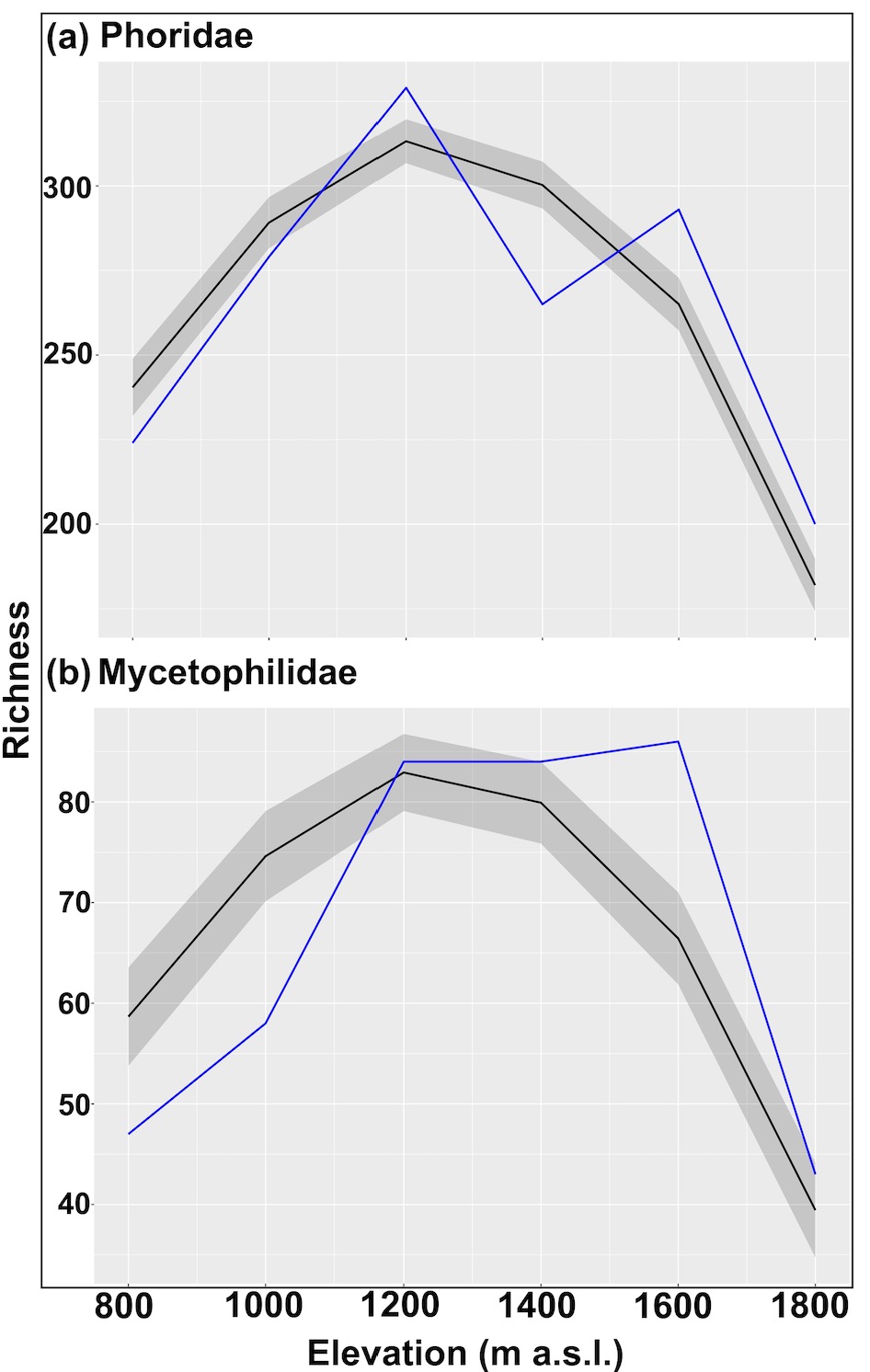

**Supplementary Figure S13.** Richness curves for (a) Phoridae and (b) Mycetophilidae in blue. Solid black lines indicate predicted values under the mid-domain effect model and the grey areas represent the expected 2.5% and 97.5% percentiles of the mOTUs richness under the MDE model.
